# Transcriptional responses to priority effects in nectar yeast

**DOI:** 10.1101/2024.01.30.578081

**Authors:** Callie R. Chappell, Pagé C. Goddard, Lexi-Ann Golden, Jonathan Hernandez, Daniela Ortiz Chavez, Marziiah Hossine, Sur Herrera Paredes, Tadashi Fukami, Manpreet K. Dhami

**Affiliations:** Department of Biology, Stanford University, Stanford, CA 94305, USA; Department of Genetics, Stanford University, Stanford, CA 94305, USA; University of Michigan Medical School, Ann Arbor, MI 48109, USA; Biochemistry and Molecular Genetics, University of Colorado Anschutz Medical Campus, Aurora, CO 80045, USA; Cellular and Molecular Biology, California State University, Northridge, Northridge, CA 91330; International Laboratory for Human Genome Research, Universidad Nacional Autónoma de México (UNAM), Juriquilla Querétaro, Querétaro, 76230, México; Department of Earth System Science, Stanford University, Stanford, CA 94305, USA; Biocontrol and Molecular Ecology, Manaaki Whenua - Landcare Research, Lincoln 7608, New Zealand; School of Biological Sciences, University of Auckland, 1142, New Zealand

**Author notes:** **Corresponding author** Callie R. Chappell.

**Keywords:** ecological genomics, niche preemption, eQTL, population genomics

## Abstract

Priority effects, where the order and timing of species arrival influence the assembly of ecological communities, have been observed in a variety of taxa and habitats. However, the genetic and molecular basis of priority effects remains unclear in most cases, hindering the mechanistic understanding of priority effects. We sought to gain such understanding for the common nectar yeast Metschnikowia *reukaufii*, which engages in strong priority effects with other species of nectar yeast, including *M. rancensis*, another species commonly found in the nectar of our study plant, the hummingbird-pollinated *Diplacus* (*Mimulus*) *aurantiacus*. After inoculation into two contrasting types of synthetic nectar simulating early arrival of *M. rancensis*, we conducted whole-transcriptome sequencing of 108 genetically diverse strains of *M. reukaufii*. We found that several genes were differentially expressed in *M. reukaufii* strains when the nectar had been conditioned by growth of *M. rancensis*. Many of these genes were associated with amino acid metabolism, consistent with our previous finding that early-arriving species limit late-arriving species’ growth by reducing amino acid availability. Furthermore, investigation of expression quantitative trait loci (eQTLs) revealed that genes involved in amino acid transport and resistance to antifungal compounds were enriched in genetic variants, with differing effects on gene expression based on priority effects of *M. rancensis*. These results demonstrate that intraspecific variation in the ability of nectar yeasts to respond to nutrient limitation and direct fungal competition may underpin the molecular mechanisms of priority effects.

## 1 Introduction

One particularly challenging type of interspecific interactions to predict are those that are historically contingent, where the outcome depends on the order and timing of species arrival (Drake, 1991; Fukami, 2015; Palmgren, 1926). These interactions, called priority effects, can be a major source of unexplained variation in community composition and function (Delory et al., 2019; Slatkin, 1974; Song et al., 2021), yet the mechanisms that cause priority effects are often unknown.

The strength of priority effects can vary depending on the genotypic identity of the species involved (Urban & De Meester, 2009). Although such interspecific variation makes priority effects more difficult to understand, this variation at the same time provides a largely unexploited opportunity to study the genetic mechanisms of priority effects. For example, our previous work with the nectar yeasts isolated from the hummingbird-pollinated shrub *Diplacus* (*Mimulus*) *aurantiacus* in California has suggested that priority effects on the dominant yeast in this system, *Mestchnikowia reukaufii*, differ in strength due to intraspecific genetic variation (Chappell et al., 2022; Dhami et al., 2016, 2018). However, specific genes responsible for this variation remain unknown.

Here, we study how intraspecific genetic variation within *M. reukaufii* may alter their susceptibility to priority effects using population transcriptomics. Standing genetic diversity may alter a species’ ability to adapt or respond to priority effects by altering transcriptional responses (Pai, Pritchard & Gilad 2015) due, for example, to differences in transcriptional regulation (Lema et al., 2019). Comparing intraspecific genetic variation and transcriptional variation can yield insight into the molecular mechanisms that explain ecological interactions such as priority effects. We studied the gene expression of 108 genome-resequenced strains of *M. reukaufii* in response to a competitor in order to identify potential molecular mechanisms of priority effects.

## 2 Materials and Methods

### 2.1 Microbial growth and sampling

A total of 108 isolates of *M. reukaufii* reported in Dhami *et al*. (2018) (Figure 1A,B, Supplementary Table S1) were plated from glycerol stocks that had been kept at -80°C on yeast mold (YM) agar and incubated at 25°C for 2 days. A single isolated colony on each of these plates was inoculated into YM broth and shaken at 200 RPM for 2 days at 25°C. Cells were pelleted at 3000 RPM for 5 minutes and washed twice with sterile phosphate buffered saline (PBS). After measuring optical density at 600 nm (OD_600_), cells were inoculated at a final density of 2×10^2^ cells/µL into 3 mL of synthetic nectar into the two nectar treatments, either the high amino acid nectar or *M. rancensis*-conditioned nectar. Four replicates per treatment were shaken at 200 RPM for 10 hours at 25°C and harvested by pelleting at 4000 RPM for 5 minutes at 25°C. The pellet was flash frozen in liquid nitrogen for subsequent RNA extraction. All 864 samples were grown and processed in a single batch over a single 48 hour period. Samples were stored at -80°C until further use.

We prepared synthetic nectar to mimic *D. aurantiacus* nectar in amino acid and sugar composition (Peay et al., 2012). Specifically, the synthetic nectar contained fructose (4%), glucose (2%), sucrose (20%), serine (0.102 mM), glycine (0.097 mM), proline (0.038 mM), glutamate (0.035 mM), aspartic acid (0.026 mM), GABA1 (0.023 mM), and alanine (0.021 mM). All components were mixed in water until dissolved, filtered through a 0.2 µm filter, and stored at –20°C until used. The nectars used in this experiment were modified in one of two ways. The first treatment increased the amino acid concentration in nectar to 10X the concentration of the standard nectar, to mimic a rich environment. The second treatment was prepared exactly the same as the first treatment, except that it was conditioned by *Metschnikowia rancensis*, which is closely related to *M. reukaufii* (Peay et al., 2012; Pozo et al., 2016) and has previously been shown to engage in strong priority effects with *M. reukaufii* (Dhami et al., 2016, 2018; Grainger et al., 2019; Peay et al., 2012; Vannette & Fukami, 2014). This *M. rancensis*-conditioned nectar was prepared by growing *M. rancensis* in the normal synthetic nectar for two days, then removing the *M. rancensis* by filtration through a 0.02 µm filter. No *M. rancensis* RNA was detectable in the final conditioned nectar.

### 2.2 RNA extraction, library preparation, and sequencing

RNA was extracted from cell pellets using a phenol-chloroform extraction method (Collart & Oliviero, 1993; Eggermont et al., 1996). Pellets were resuspended in phenol, chloroform, and isoamyl alcohol with an RNA extraction buffer, and incubated at 60°C for 30 minutes while shaking. RNA was extracted using phase separation and centrifugation with phenol, chloroform, and isoamyl alcohol. RNA was precipitated using a lithium chloride solution and incubated at -20°C overnight. Samples were centrifuged, and the pellet resuspended in cold 80% ethanol. After pelleting again, the ethanol was removed and the purified RNA was resuspended in pyrogen-free water and stored at -80°C. Diluted RNA was treated with DNase I using a Zymo DNase I kit (Zymo Research, Irvine, CA) and purified using a Zymo RNA Clean and Concentrator Kit (Zymo Research, Irvine, CA). Final RNA concentrations were quantified using an Invitrogen Quant-iT Broad Range RNA Assay Kit (Thermo Fisher, Waltham, MA). Purified RNA was shipped to the Joint Genome Institute (JGI), where cDNA libraries were generated using an TruSeq RNA Library Prep Kit (Illumina, San Diego, CA). Samples were sequenced on a NovaSeq 6000 sequencer (2 × 251bp reads). JGI provided cleaned, trimmed reads for bioinformatic analysis. BBDuk (version 38.25) was used to remove contaminants, trim reads of adapter sequences, and remove reads with low quality scores. BBMap was used to map reads to an *M. reukaufii* reference genome (Dhami et al., 2016) and filter reads that mapped to human, cat, dog, or mouse references at 93% identity, as well as ribosomal sequences.

### 2.3 Differential expression analysis

In R (3.5.2), DESeq2 (Love et al. 2014) was used to generate a differential expression count matrix, using a raw read count matrix as the input. The transformed count matrix was used in principal component analysis (PCA) and differences between groups were analyzed by PERMANOVA with Bray-Curtis Dissimilarity using the adonis function in vegan (2.5-7) (Oksanen et al., 2019). After log normalization, independent hypothesis weighting was used to determine the number of significantly differentially expressed genes per treatment (p<0.1, FDR=0.1). Significantly differentially expressed genes (p<0.05, LFC < 0.5 or >0.5) are reported in Supplementary Table S2. We predicted the function of expressed genes using the InterPro database (interproscan *5*.*45–80*.*0*) (Blum et al., 2021). TopGO (2.46.0) was used to identify enriched GO terms in significantly differentially expressed genes (Alexa & Rahnenführer, 2009). GO term enrichment was examined for Biological Process (BP) (Figure 2B), Cellular Component (CC), and Molecular Function (MF) based on the “weight01” algorithm and Fisher statistic (Supplementary Table S3).

### 2.4 cis-eQTL Mapping

Pre- and post-processing of the cis-QTL data were performed in R (4.0.3). Strains with no genotype data, and samples with fewer than 4,000,000 RNA-seq library reads were excluded from eQTL calling. Samples corresponding to the Y76 strain had exceptionally low read counts relative to the rest of the dataset (mean = 1,557,935 vs 10,582,904 reads). A minimum library size of four million reads provided an arbitrary cutoff that cleanly separated these low-expression samples from the rest.

Only genes with at least 0.1 TPM (transcript per kilobase million) and five or more read counts in at least 144 samples (20% of samples) were retained for analysis. Genes on scaffolds without complete genotype information were excluded from cis-eQTL mapping. Raw gene count estimates were converted to counts per million (cpm) using the cpm function from edgeR (Robinson et al., 2010), then inverse-normalized using the RankNorm function from RNOmni (McCaw et al., 2020). Principal components (PCs) were calculated from the transformed count data using PCATools (Blighe, 2018/2022), and the number of PCs to include in the final model was determined using the Gavish-Donoho method (Gavish & Donoho, 2014).

eQTL mapping was performed using tensorQTL in Python (3.9.8) (Taylor-Weiner et al., 2019). Eight samples representing the Y76 strain were excluded from analysis due to low library size, an additional 32 samples representing the Y413, Y962, and Y983 strains were excluded due to a lack of genotype data, and 77 samples across 44 strains were removed from analysis due to lack of RNA-seq data, leaving 718 remaining samples across both conditions for eQTL calling. Lack of genotype information for scaffolds 117 and 100 led to the exclusion of 1398 and 2 genes, respectively. Of the remaining 4589 annotated genes, 252 genes were removed from analysis due to low expression, resulting in 4337 (94.5%) genes for eQTL analysis. No variants were dropped. Variants within 1 Mb of the transcription start site were tested for association with gene expression using a linear model with an interaction term. The model corrected for sample strain, genotype sub-population, collection site, sequencing plate, and the first 33 PCs:

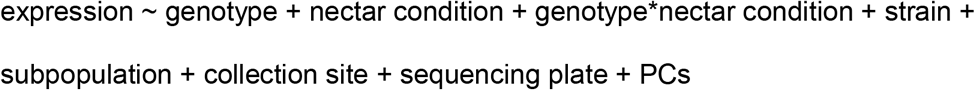

The top eQTL variant was reported for each gene, and multiple testing correction by multiplying the nominal p-values effective number of tests reported by *tensorQTL*’s eigenMT option.

Response eQTL genes were defined by a significant interaction term (Bonferroni corrected adjusted p-value <= 0.05). Downstream assessments were carried out in R (4.0.3).

We predicted the function of expressed genes using the InterPro database (`interproscan 5.45–80.0`) [4]. TopGO (2.46.0) was used to identify enriched GO terms in genes associated with interaction eQTLs [9]. GO term enrichment was examined for Biological Process (BP) based on the “weight01” algorithm and Fisher statistic.

## 3 Results

### 3.1 Transcriptome sequencing and alignment

A total of 12,559,122,137 raw reads were obtained from all samples. After quality control, an average of 14,428,231 (85%) of clean reads per sample mapped to the *M. reukaufii* MR1 reference genome (Dhami et al., 2016) (Supplementary Table S1, Supplementary Table S2, Figure S1). Distributions of mapped reads were similar between the two nectar-type treatments.

### 3.2 Population-level response to the nectar environment

Samples were clustered by the three groupings we previously described (Dhami et al., 2018) (Figure 1C) (PERMANOVA, population group: n=718, R^2^=0.028, p=0.009). We refer to the groups described in Dhami et al. (2018) as populations in this paper, following the convention in population genomics. Thus, by populations, we do not necessarily mean spatially defined entities. In addition, we found two clusters that consisted of strains belonging to population 1 (Figure 1C), which corresponded to the two genetically distinct subgroups of population 1 (Figure 1B). There was no statistically significant difference between the nectar treatments (PERMANOVA, nectar treatment: n=718, R^2^=0.0007, p=0.60).

**Figure 1:**
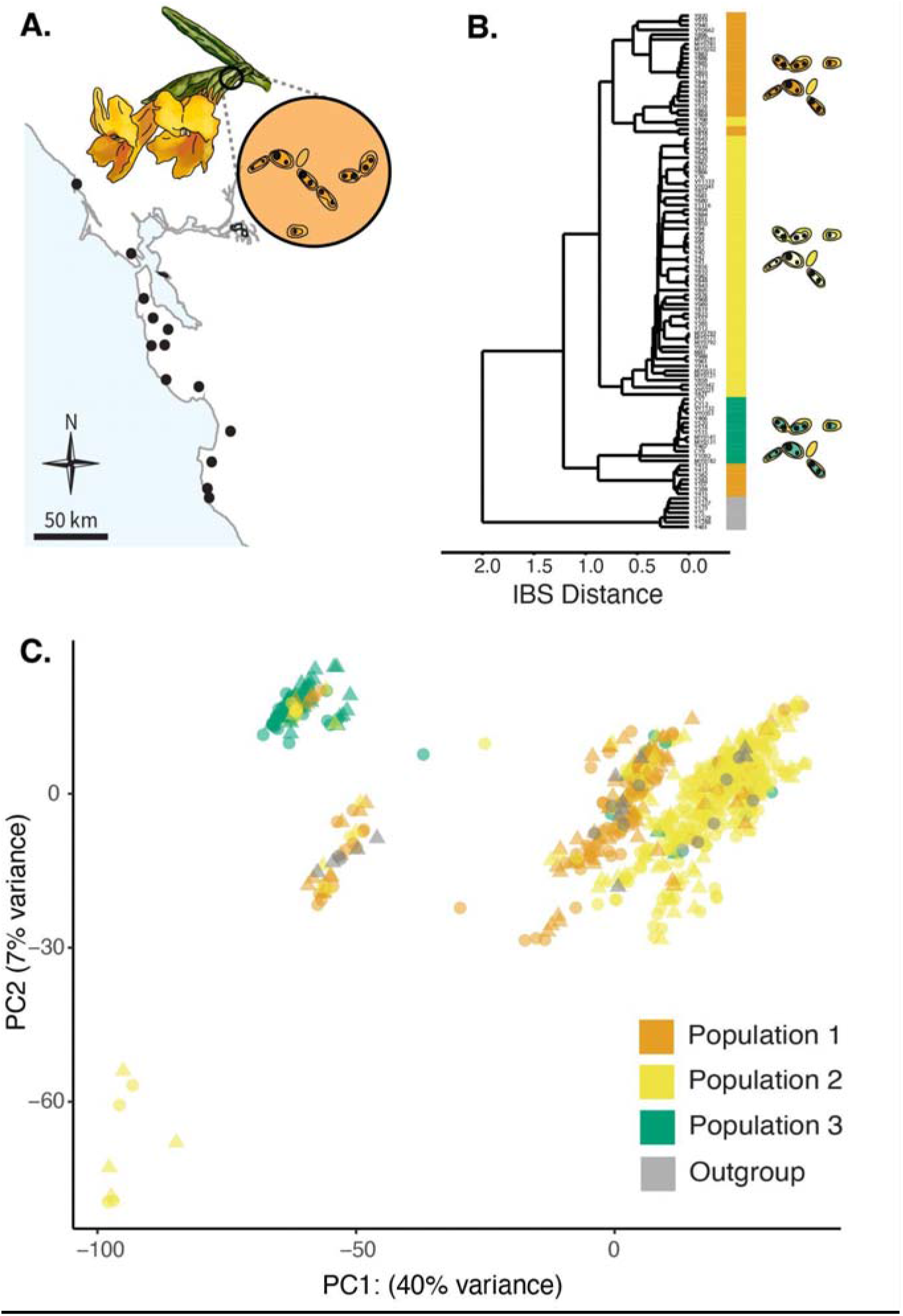
Sub-populations of nectar yeast differ in transcriptional response. (A) Wild yeast were isolated from the nectar of *Diplacus aurantiacus* growing at 12 sites across the Greater Bay Area of California. (B) Whole genome resequencing of 108 strains of nectar yeast *Metschnikowia reukaufii* revealed three populations (Dhami et al., 2018). (C) In the current study, 108 strains of *M. reukaufii* were grown separately in synthetic *D. aurantiacus* nectar with high N or synthetic nectar conditioned with competitor yeast *M. rancensis*. After 10 hours of growth, overall gene expression was predominantly influenced by population group (color) and less by nectar treatment (*M. reukaufii* grown in high N nectar are circles vs. conditioned nectar as triangles) (PERMANOVA, population group: n=742, R^2^=0.028,, p=0.009). There was no significant difference between nectar treatments (PERMANOVA, nectar treatment: n=718, R^2^=0.0007, p=0.60)

### 3.3 Functional enrichment of differentially expressed genes

A total of 4,972 genes had a nonzero total read count when mapped to the reference *M. reukaufii* MR1 genome (Dhami et al., 2018) (Figure S2, S3). Of these, 1,177 (20%) genes were more highly expressed in the high N nectar, and 886 (15%) were more highly expressed in the conditioned nectar after accounting for independent hypothesis weighting (p adj. <0.1, nominal FDR control=0.1) (Figure 2A, Supplementary Table S3). Of differentially expressed genes, 1,657 (90%) had predicted functions. Biological processes enriched in differentially expressed genes by treatment included cellular nitrogen biosynthesis (n=193, p=0.01) and organonitrogen compound biosynthesis (n=165, p=0.08) (Figure 2B, Supplementary Table S4, Supplementary Figure S4).

**Figure 2:**
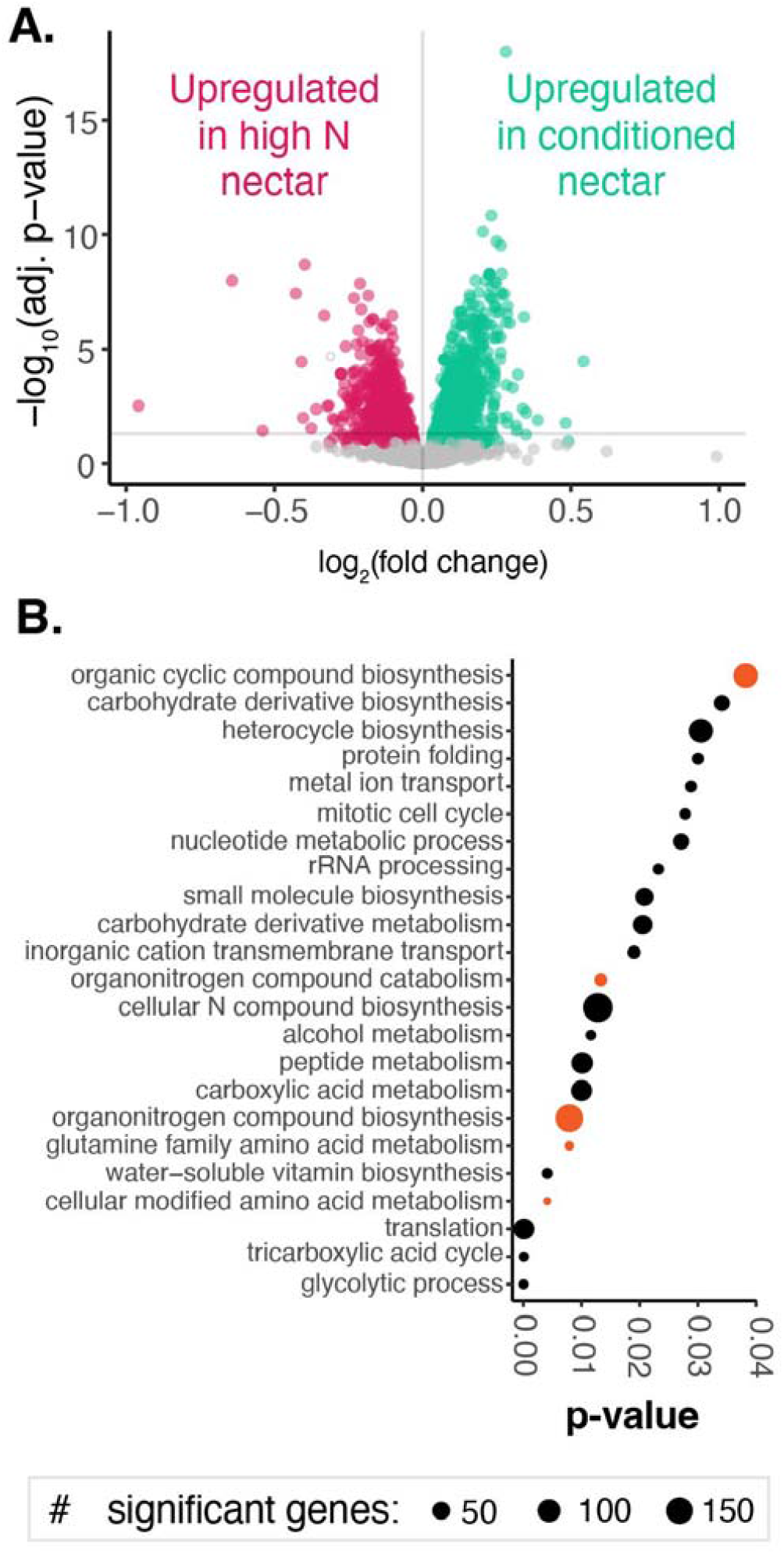
Transcriptional differences based in nectar treatment. (A) Significantly differentially expressed genes (n=6,613) between treatments, high N nectar (left, magenta) and *M. rancensis-*conditioned nectar (right, teal). (B) Enriched biological process gene ontology (GO) terms in significantly differentially expressed genes (n=2,520). The size of each bubble denotes the number of significant genes with that GO term. GO terms colored in orange are associated with amino acid metabolism or catabolism.

### 3.4 eQTLs

eQTL mapping was performed using 88,193 SNPs and 4,111 genes, testing the association between genotype and transcriptional response to the two nectar treatments. We observed 355 genes whose expression was associated with genotype (genotype eQTLs), and 176 with treatment-dependent genotype associations (interaction eQTLs, Figure 3A). Only 39 genes were shared between genotype and interaction eQTLs. The effect sizes of significant eQTL were similar between treatment, genotype, and the interaction between them (Figure 3B).

**Figure 3:**
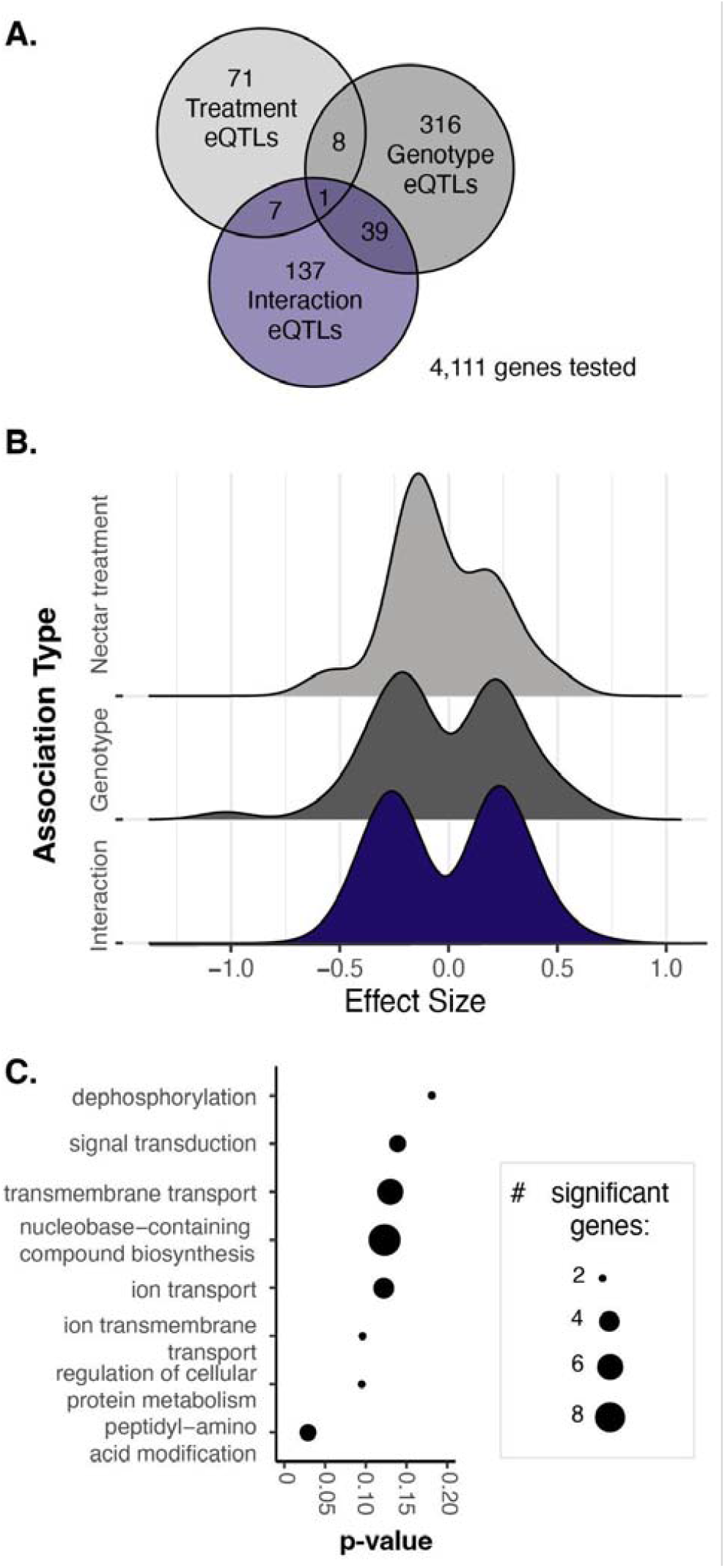
Expression QTLs by treatment, genotype, and interaction. (A) Summary of eQTLs by treatment, genotype, and the interaction between them. (B) Frequency distribution of effect sizes by treatment, genotype, and interaction. (C) GO term enrichment of interaction eQTLs.

Generally, treatment, genotype, and interaction eQTLs had the same direction of effect (Supplementary figure S5).

The top GO terms of genes associated with interaction eQTLs included ion transport pathways (Figure 3C). For example, a predicted aminotransferase involved in the N starvation response was affected by treatment (Figure 4A) (adj. p_GxE_=0.013), whereas an ATP-binding cassette (ABC) transporter was differentially affected by genotype and nectar treatment (adj. p_GxE_=0.002) (Figure 4B) (Supplementary table S5).

**Figure 4:**
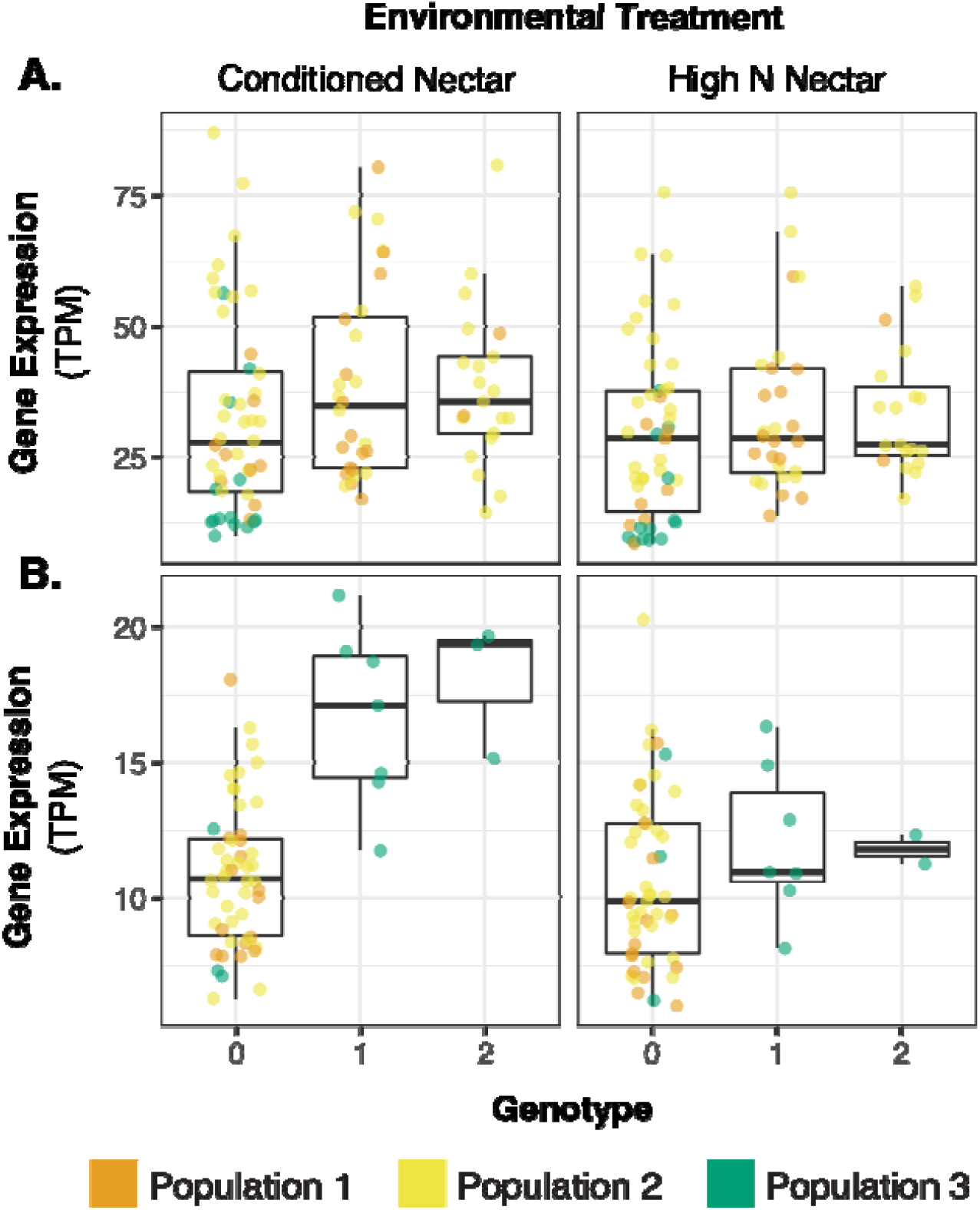
Differential expression by genotype and treatment. (A) Expression (transcript per kilobase million, or TPM) of a putative amino acid transporter (transcript 38411) based on genotype at SNP 764027 and nectar treatment (adj. P_GxE_=0.013, effect size=0.100+/-0.022). (B) Expression (TPM) of a putative ABC transporter (transcript 35539) based on genotype at SNP 1047038 and nectar treatment (adj. P_GxE_=0.002, effect size=-0.370+/-0.075). Individual points represent gene expression within an individual sample. Points are colored by whether the sample (strain) was from population 1, 2, or 3 (Figure 4A).

## 4 Discussion

Functional analysis of genes that were differentially expressed between the two nectar environments found enrichment associated with cellular nitrogen biosynthesis and organonitrogen compound biosynthesis (Figure 2B). In nectar, competition for amino acids between *M. reukaufii* and other yeast species such as *M. rancensis* can be severe (Dhami et al., 2018; Grainger et al., 2019; Letten et al., 2018; Peay et al., 2012; Vannette & Fukami, 2014). When *M. rancensis* arrives first to a *D. aurantiacus* flower via a hummingbird, *M. rancensis* can deplete amino acids in nectar. Thus, to resist priority effects by *M. rancensis*, a late-arriving *M. reukaufii* strain would have to express genes that help them survive in low N environments. Indeed, our differential expression analysis shows that *M. reukaufii* changes expression of genes associated with competition for amino acids due to the early arrival of *M. rancensis*, suggesting this competition as a mechanism of priority effects. This finding is broadly consistent with our prior genome analysis of *M. reukaufii*, which revealed extensive gene duplications in high-capacity amino acid transporters (Dhami et al., 2016). We speculate that the limited nitrogen availability in the form of amino acids, an essential N resource for microbial growth, have exerted strong selection pressure in the evolution of *M. reukaufii* (Dhami et al. 2018).

A surprising finding from this study is how strongly genotypic identity influences the yeasts’ molecular response to nectar conditions. Prior work identified three groups within this collection of *M. reukaufii* strains (Dhami et al., 2018) (Figure 1B). Our results suggest that the three groups do not always respond in the same way to priority effects. By combining population-level genetic variation (Figure 1) and differences in gene expression (Figure 2), we identified several genes whose expression level differed based on yeast lineage (*cis* eQTLs) (Figure 3).

Two examples of eQTLs illustrate how our results suggest lineage-specific mechanisms of priority effects. First, variation at SNP 764027 demonstrates how the interaction between genotype and the environment can strongly alter expression of a putative amino acid transporter (transcript 38411) (Figure 4A). This gene, which is predicted to help import amino acids into the cell, was more highly expressed in nectar depleted of amino acids by *M. rancensis*. Increased expression of a putative amino acid import gene in nectar environment conditioned by a competitor suggests niche preemption (sensu Fukami, 2015) as a mechanism of priority effects. In the conditioned nectar treatment, early arriving *M. rancensis* probably drew down nectar N, making the late-arriving *M. reukaufii* express amino acid transporters to scavenge for the remaining amino acids. However, our data indicate that the extent to which *M. reukaufii* can do so depends on the genotype: strains homozygous for the alt allele at SNP 764027 are able to express that gene more highly than those with either alternative allele.

A second example shows how the genotype effects can be reversed, depending on the environmental treatment. In the conditioned nectar treatment, there was increased expression of a transcript (35539) depending on the genotype at SNP 1047038. However, the effect of genotype was reversed in the high-N nectar environment (Figure 4B). This gene is predicted to be an ABC transporter, which can support resistance to antifungal compounds (Kumari et al., 2021). In the nectar conditioned by *M. rancensis*, potentially containing antifungal compounds that inhibit *M. reukaufii* growth (Álvarez-Pérez et al., 2016), *M. reukaufii’s* expression of this ABC transporter differed based on its genotype at SNP 1047038. This second example also supports a niche preemption mechanism of priority effects, where *M. rancensis* could prevent the growth of later-arriving *M. reukaufii* by producing a toxic compound (Álvarez-Pérez et al., 2016).

The interacting effect of environment (such as niche preemption by *M. rancensis*) and genotype underscores how the effects of intraspecific genetic variation within *M. reukaufii* can only be understood in their environmental context. The two examples just discussed, as well as the 137 additional interaction eQTLs (Figure 3A), suggest that neither yeast genotype nor nectar environment alone fully explains the molecular response to priority effects. Instead, the interplay between genotype and environment is necessary to predict the mechanisms by which species interact.

Here we have sought to connect population-level genetic variation to differences in molecular traits (gene expression) associated with response to priority effects. Future research could more directly test for the effects of the proposed genes on susceptibility to priority effects in *M. reukaufii*. For example, transgenic *M. reukaufii* with knock-out genes or eQTLs identified in this study could directly test their impact on priority effects. Studies like these that make use of interspecific genetic variation in the wild should help us with improving our understanding of the genetic basis of priority effects. Genetic tools such as population transcriptomics can uncover specific mechanisms by which species interact.

## Data Accessibility and Benefit-Sharing

### Genetic data

Raw sequence reads are downloadable from the JGI Genome Portal (FD 1202157; SP 1202175; AP 1202158).

Individual genotype data, sample metadata, and analyses are available on GitLab: https://gitlab.com/teamnectarmicrobe/n01_nectar_yeast_transcriptomics

### Sample metadata

Metadata are also stored on GitLab: https://gitlab.com/teamnectarmicrobe/n01_nectar_yeast_transcriptomics
**Benefits Generated:** This research occurred in collaboration with several undergraduate researchers, whose training in ecology, evolution, and genomics was advanced through this work. Broadly, our group is committed to advancing diversity, equity, and inclusion in the life sciences.

## Author Contributions

CRC, MKD, and TF designed and conducted the research. CRC, PCG, LG, JH, DOC, MH, and SHP analyzed the data. CRC and TF wrote the paper, and all authors contributed to improving the paper.

## Acknowledgements

We thank Laura Bogar, Ben Bowen, Glade Dlott, Adina Howe, Grant Kinsler, Katherine Louie, Trent Northen, Lauren O’Connell, Kabir Peay, Dmitri Petrov, Gavin Sherlock, Rachel Vannette, Tuya Yokoyama, and the members of the community ecology group at Stanford for discussion and comments. Funding: This work was supported by the National Science Foundation (DEB 1149600, DEB 1737758), Community Science Program (CSP) grant from the Joint Genome Institute, and Stanford University’s Terman Fellowship. CRC was supported by a National Science Foundation Graduate Research Fellowship (DGE 1656518) and a Stanford Graduate Fellowship. MKD was supported by Marsden Fund Grant (MFP-LCR-2002). SHP was supported by the Life Sciences Research Foundation. LG and JH were supported by the Stanford Department of Biology VPUE Biology Summer Research Program. DOC and MH were supported by the Stanford Summer Research Program.

## References

Alexa, A., & Rahnenführer, J. (2009). Gene set enrichment analysis with topGO. Bioconductor Improv, 27, 1–26.

Álvarez-Pérez, S., de Vega, C., Pozo, M. I., Lenaerts, M., Van Assche, A., Herrera, C. M., Jacquemyn, H., & Lievens, B. (2016). Nectar yeasts of the Metschnikowia clade are highly susceptible to azole antifungals widely used in medicine and agriculture. FEMS Yeast Research, 16(1). 10.1093/femsyr/fov115

Blighe, K. (2022). PCAtools: Everything Principal Component Analysis [R package version 2.10.0]. https://github.com/kevinblighe/PCAtools (Original work published 2018)

Blum, M., Chang, H.-Y., Chuguransky, S., Grego, T., Kandasaamy, S., Mitchell, A., Nuka, G., Paysan-Lafosse, T., Qureshi, M., Raj, S., Richardson, L., Salazar, G. A., Williams, L., Bork, P., Bridge, A., Gough, J., Haft, D. H., Letunic, I., Marchler-Bauer, A., … Finn, R. D. (2021). The InterPro protein families and domains database: 20 years on. Nucleic Acids Research, 49(D1), D344–D354. 10.1093/nar/gkaa977

Chappell, C. R., Dhami, M. K., Bitter, M. C., Czech, L., Herrera Paredes, S., Barrie, F. B., Calderón, Y., Eritano, K., Golden, L.-A., Hekmat-Scafe, D., Hsu, V., Kieschnick, C., Malladi, S., Rush, N., & Fukami, T. (2022). Wide-ranging consequences of priority effects governed by an overarching factor. eLife, 11, e79647. 10.7554/eLife.79647

Collart, M. A., & Oliviero, S. (1993). Preparation of yeast RNA. Current Protocols in Molecular Biology, 13.12.1–13.12.5.

Delory, B. M., Weidlich, E. W. A., von Gillhaussen, P., & Temperton, V. M. (2019). When history matters: The overlooked role of priority effects in grassland overyielding. Functional Ecology, 33(12), 2369–2380. 10.1111/1365-2435.13455

Dhami, M. K., Hartwig, T., & Fukami, T. (2016). Genetic basis of priority effects: Insights from nectar yeast. Proceedings of the Royal Society of London B: Biological Sciences, 283(1840), 20161455. 10.1098/rspb.2016.1455

Dhami, M. K., Hartwig, T., Letten, A. D., Banf, M., & Fukami, T. (2018). Genomic diversity of a nectar yeast clusters into metabolically, but not geographically, distinct lineages. Molecular Ecology, 27(8). 10.1111/mec.14535

Drake, J. A. (1991). Community-assembly mechanics and the structure of an experimental species ensemble. The American Naturalist, 137(1), 1–26. https://www.jstor.org/stable/2462154

Eggermont, K., Goderis, I. J., & Broekaert, W. F. (1996). High-throughput RNA extraction from plant samples based on homogenisation by reciprocal shaking in the presence of a mixture of sand and glass beads. Plant Molecular Biology Reporter, 14(3), 273–279.

Fukami, T. (2015). Historical contingency in community assembly: Integrating niches, species pools, and priority effects. Annual Review of Ecology, Evolution, and Systematics, 46(1), 1–23. 10.1146/annurev-ecolsys-110411-160340

Gavish, M., & Donoho, D. L. (2014). The Optimal Hard Threshold for Singular Values is \(4/\sqrt {3}\). IEEE Transactions on Information Theory, 60(8), 5040–5053. 10.1109/TIT.2014.2323359

Grainger, T. N., Letten, A. D., Gilbert, B., & Fukami, T. (2019). Applying modern coexistence theory to priority effects. Proceedings of the National Academy of Sciences, 116(13), 6205–6210. 10.1073/pnas.1803122116

Kumari, S., Kumar, M., Gaur, N. A., & Prasad, R. (2021). Multiple roles of ABC transporters in yeast. Fungal Genetics and Biology, 150, 103550. 10.1016/j.fgb.2021.103550

Lema, K. A., Metegnier, G., Quéré, J., Latimier, M., Youenou, A., Lambert, C., Fauchot, J., & Le Gac, M. (2019). Inter- and Intra-Specific Transcriptional and Phenotypic Responses of Pseudo-nitzschia under Different Nutrient Conditions. Genome Biology and Evolution, 11(3), 731–747. 10.1093/gbe/evz030

Letten, A. D., Dhami, M. K., Ke, P.-J., & Fukami, T. (2018). Species coexistence through simultaneous fluctuation-dependent mechanisms. Proceedings of the National Academy of Sciences, 115(26), 6745–6750. 10.1073/pnas.1801846115

McCaw, Z. R., Lane, J. M., Saxena, R., Redline, S., & Lin, X. (2020). Operating characteristics of the rank-based inverse normal transformation for quantitative trait analysis in genome-wide association studies. Biometrics, 76(4), 1262–1272. 10.1111/biom.13214

Oksanen, J., Blanchet, F. G., Friendly, M., Kindt, R., Legendre, P., McGlinn, D., Minchin, P. R., O’Hara, R. B., Simpson, G. L., Solymos, P., Stevens, M. H. H., Szoecs, E., & Wagner, H. (2019). vegan: Community Ecology Package (2.5-5) [Computer software]. https://CRAN.R-project.org/package=vegan

Palmgren, A. (1926). Chance as an element in plant geography. In B. M. Duggar (Ed.), Proceedings of the International Congress of Plant Sciences (Vol. 1, pp. 591–602).

Peay, K. G., Belisle, M., & Fukami, T. (2012). Phylogenetic relatedness predicts priority effects in nectar yeast communities. Proceedings of the Royal Society of London B: Biological Sciences, 279(1729), 749–758. 10.1098/rspb.2011.1230

Pozo, M. I., Herrera, C. M., Lachance, M.-A., Verstrepen, K., Lievens, B., & Jacquemyn, H. (2016). Species coexistence in simple microbial communities: Unravelling the phenotypic landscape of co-occurring Metschnikowia species in floral nectar. Environmental Microbiology, 18(6), 1850–1862. http://onlinelibrary.wiley.com/doi/10.1111/1462-2920.13037/full

Robinson, M. D., McCarthy, D. J., & Smyth, G. K. (2010). edgeR: A Bioconductor package for differential expression analysis of digital gene expression data. Bioinformatics, 26(1), 139–140. 10.1093/bioinformatics/btp616

Slatkin, M. (1974). Competition and Regional Coexistence. Ecology, 55(1), 128–134. JSTOR. 10.2307/1934625

Song, C., Fukami, T., & Saavedra, S. (2021). Untangling the complexity of priority effects in multispecies communities. Ecology Letters, 24(11), 2301–2313. 10.1111/ele.13870

Taylor-Weiner, A., Aguet, F., Haradhvala, N. J., Gosai, S., Anand, S., Kim, J., Ardlie, K., Van Allen, E. M., & Getz, G. (2019). Scaling computational genomics to millions of individuals with GPUs. Genome Biology, 20(1), 228. 10.1186/s13059-019-1836-7

Urban, M. C., & De Meester, L. (2009). Community monopolization: Local adaptation enhances priority effects in an evolving metacommunity. Proceedings of the Royal Society of London B: Biological Sciences, 276(1676), 4129–4138. 10.1098/rspb.2009.1382

Vannette, R. L., & Fukami, T. (2014). Historical contingency in species interactions: Towards niche-based predictions. Ecology Letters, 17(1), 115–124. 10.1111/ele.12204

