## Supplement for "Transcriptional responses to priority effects in nectar yeast"

### **Supplementary Figures**

#### **S1: Mapped reads by population**

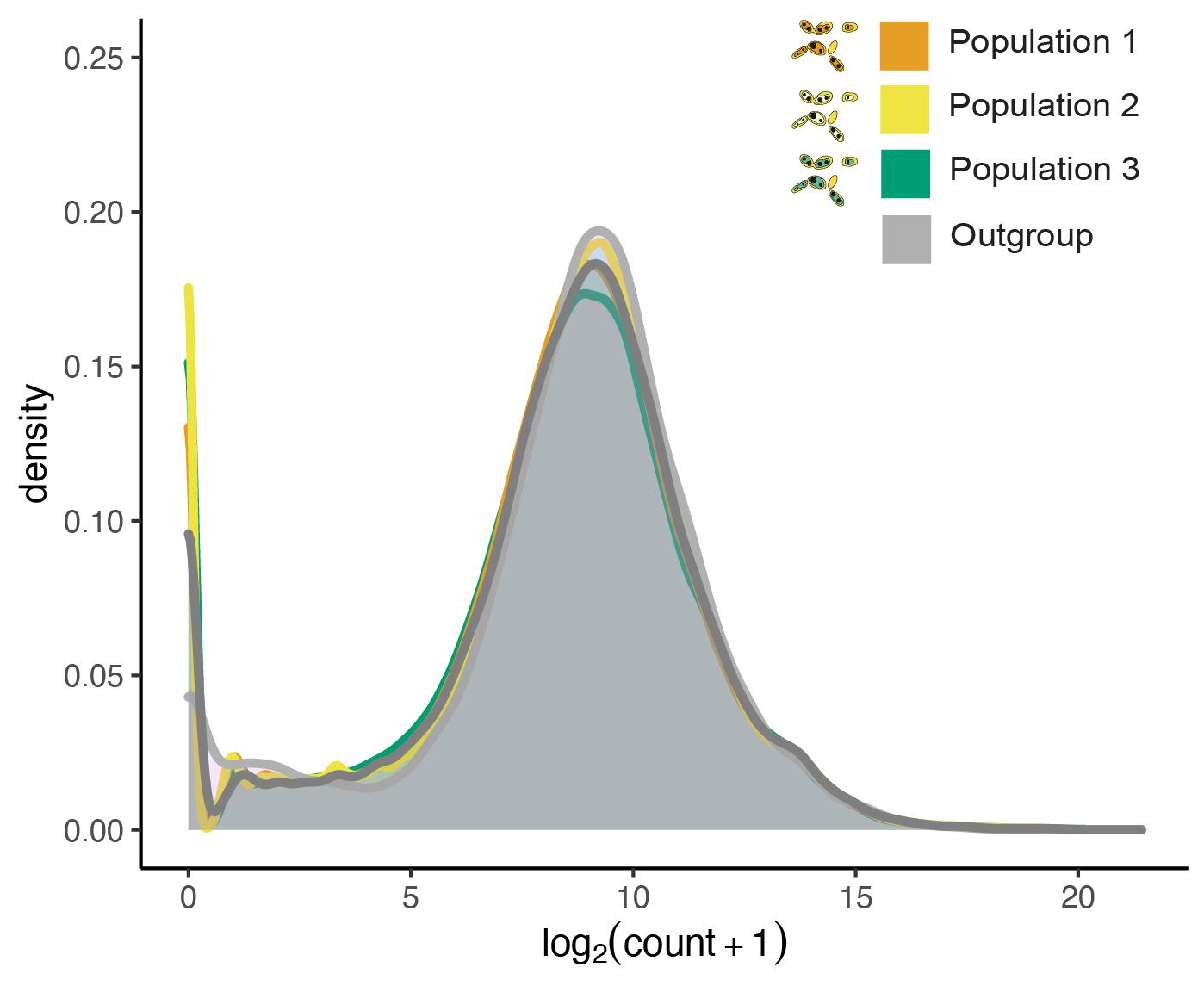

Distribution of mapped reads (log-transformed counts) for three population groups of *M. reukaufii* and the outgroup.

#### **S2: Heatmap of mapped genes**

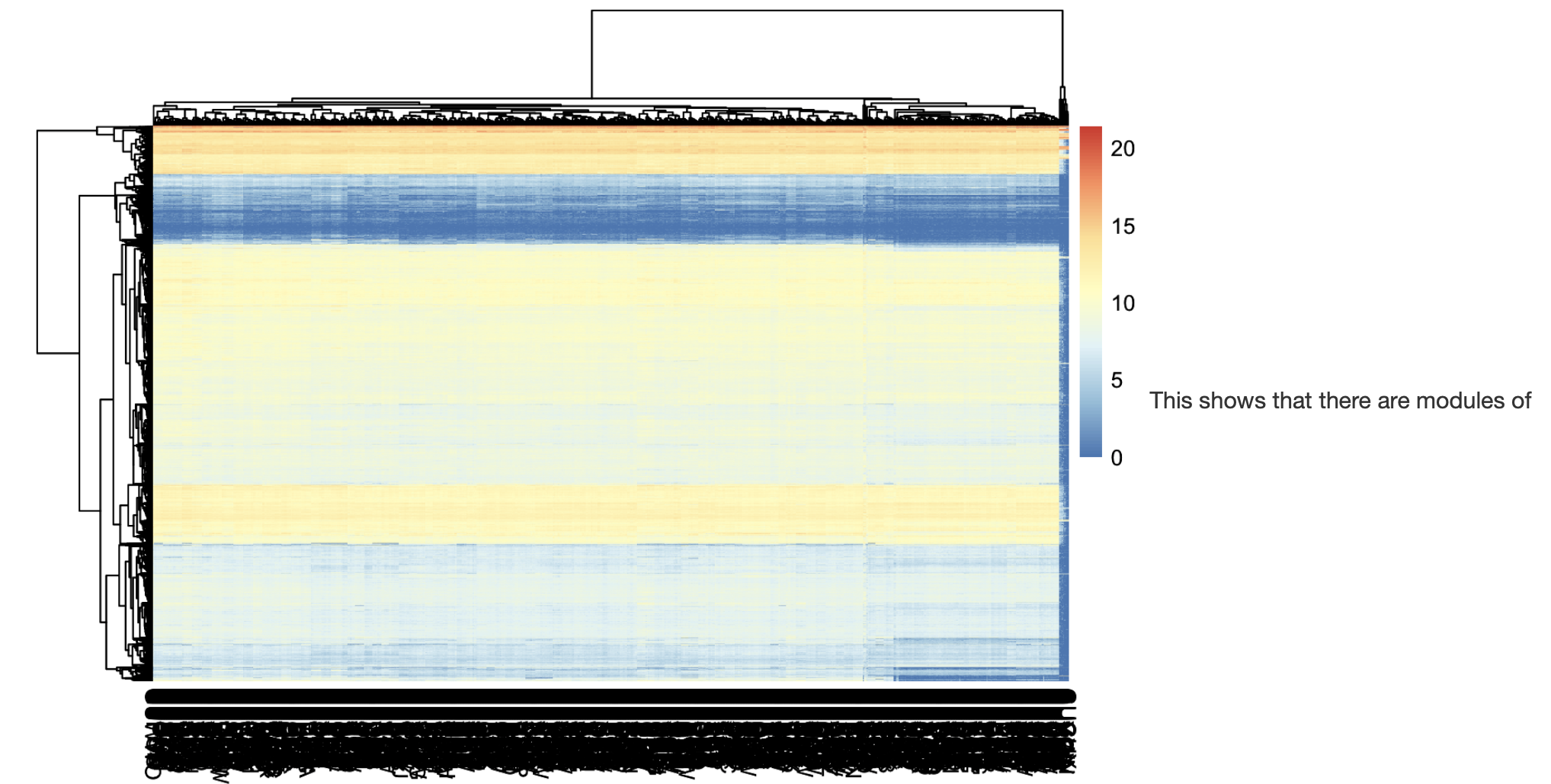

Heatmap of expression of all genes in *M. reukaufii* (rows) per sample across treatments (columns). Expression of genes was log-transformed. Red represents highly expressed genes while blue represents genes that are lowly expressed.

#### **S3: Distribution of differential expression for mapped genes by treatment**

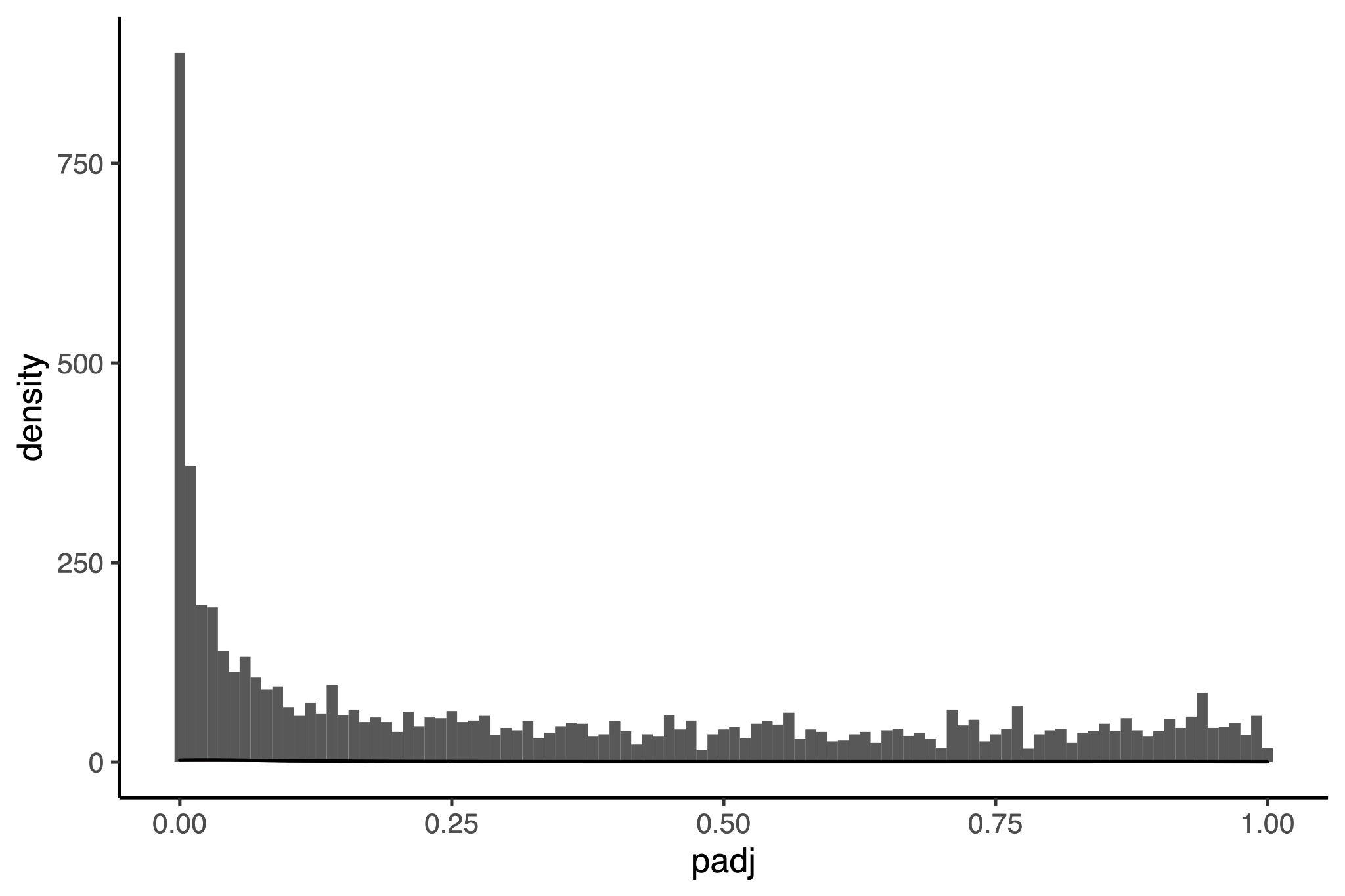

Distribution of adjusted p-values for expressed genes by treatment. There are 1846 genes with adjusted p value less than 0.05.

#### **S4: GO term enrichment network**

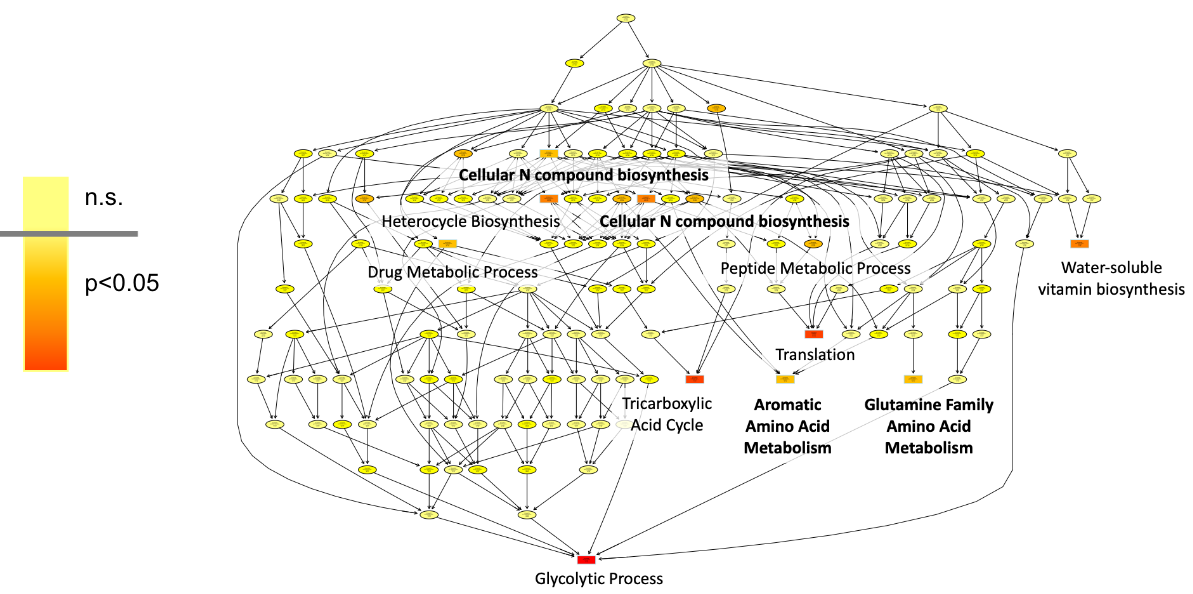

#### **S5: eQTL betas by gene**

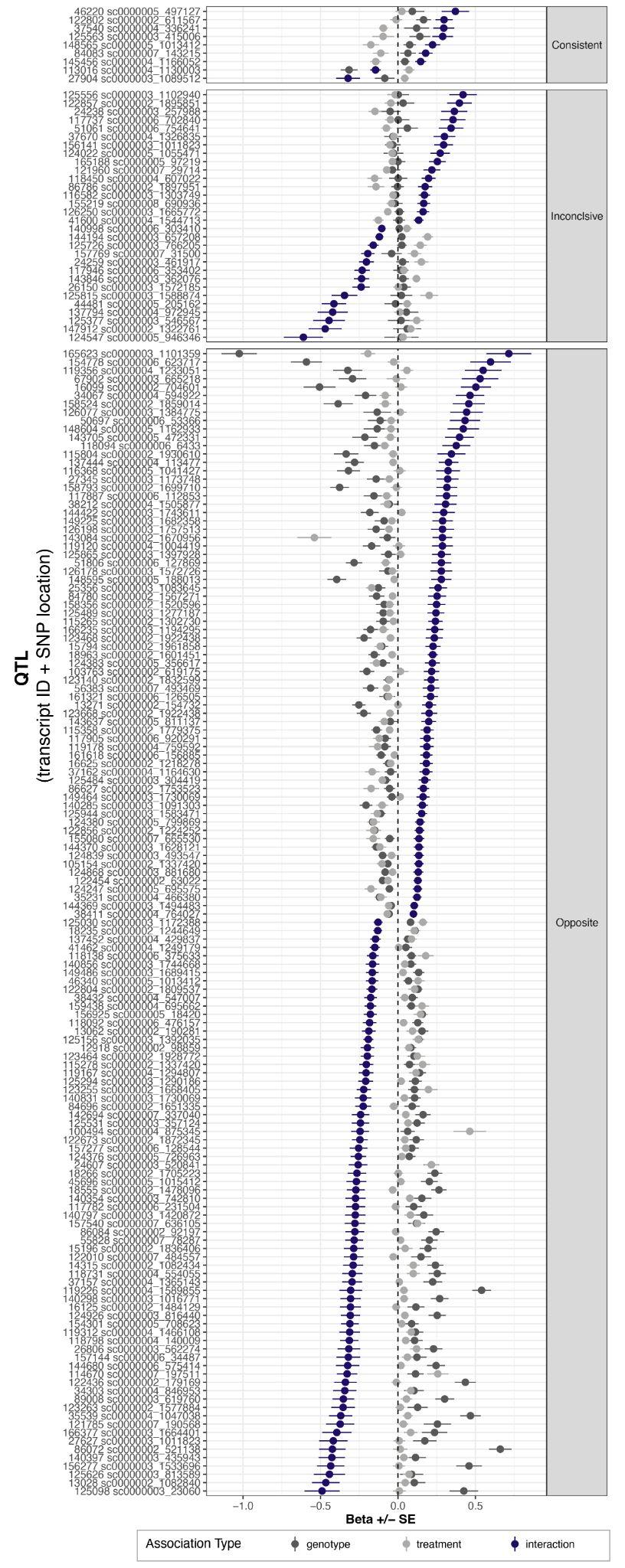

### **Supplementary Tables:**

#### **S1: *M. reukaufii strains***

| **Strain ID** | **Site** | **Isolated by** | **Collection year** | **Genotype** |
| --- | --- | --- | --- | --- |
| CY11 | Jasper Ridge | Dhami MK, et al. | 2014 | Group3 |
| CY13 | Jasper Ridge | Dhami MK, et al. | 2014 | Group1 |
| CY7 | Jasper Ridge | Dhami MK, et al. | 2014 | Group1 |
| CY9 | Jasper Ridge | Dhami MK, et al. | 2014 | Group1 |
| MR1 | Jasper Ridge | Belisle M. | 2012 | Group3 |
| MY0121 | Jasper Ridge | Dhami MK, et al. | 2014 | Group3 |
| MY0131 | Jasper Ridge | Dhami MK, et al. | 2014 | Group1 |
| MY0141 | Jasper Ridge | Dhami MK, et al. | 2014 | Group1 |
| MY0182 | Jasper Ridge | Dhami MK, et al. | 2014 | Group1 |
| MY0202 | Jasper Ridge | Dhami MK, et al. | 2014 | Group2 |
| MY0281 | Jasper Ridge | Dhami MK, et al. | 2014 | Group2 |
| MY0551 | Jasper Ridge | Dhami MK, et al. | 2014 | Group3 |
| MY0771 | Jasper Ridge | Dhami MK, et al. | 2014 | Group3 |
| MY0781 | Jasper Ridge | Dhami MK, et al. | 2014 | Group2 |
| MY0792 | Jasper Ridge | Dhami MK, et al. | 2014 | Group3 |
| MY0793 | Jasper Ridge | Dhami MK, et al. | 2014 | Group3 |
| VY0221 | Jasper Ridge | Dhami MK, et al. | 2014 | Group3 |
| VY0341 | Jasper Ridge | Dhami MK, et al. | 2014 | Group3 |
| VY0342 | Jasper Ridge | Dhami MK, et al. | 2014 | Group3 |
| VY0351 | Jasper Ridge | Dhami MK, et al. | 2014 | Group1 |
| VY0662 | Jasper Ridge | Dhami MK, et al. | 2014 | Group2 |
| VY1132 | Jasper Ridge | Dhami MK, et al. | 2014 | Group1 |
| VY1133 | Jasper Ridge | Dhami MK, et al. | 2014 | Group3 |
| Y107 | La Honda | Dhami MK, et al. | 2015 | Group2 |
| Y1092 | Swanton | Dhami MK, et al. | 2015 | Group1 |
| Y1116 | Big Sur | Dhami MK, et al. | 2015 | Group3 |
| Y1227 | Skyline Boulevard | Dhami MK, et al. | 2015 | Outgroup |
| Y1229 | Skyline Boulevard | Dhami MK, et al. | 2015 | Outgroup |
| Y1248 | Skyline Boulevard | Dhami MK, et al. | 2015 | Outgroup |
| Y173 | Swantonn Gregorio | Dhami MK, et al. | 2015 | Outgroup |
| Y174 | Swantonn Gregorio | Dhami MK, et al. | 2015 | Outgroup |
| Y312 | Muir Woods | Dhami MK, et al. | 2015 | Group3 |
| Y382 | Muir Woods | Dhami MK, et al. | 2015 | Group2 |
| Y383 | Swanton | Dhami MK, et al. | 2015 | Group2 |
| Y384 | Muir Woods | Dhami MK, et al. | 2015 | Group2 |
| Y385 | Muir Woods | Dhami MK, et al. | 2015 | Group3 |
| Y40 | La Honda | Dhami MK, et al. | 2015 | Group3 |
| Y41 | La Honda | Dhami MK, et al. | 2015 | Group3 |
| Y412 | Muir Woods | Dhami MK, et al. | 2015 | Group2 |
| Y413a | Muir Woods | Dhami MK, et al. | 2015 | Group2 |
| Y415 | Muir Woods | Dhami MK, et al. | 2015 | Group2 |
| Y42 | La Honda | Dhami MK, et al. | 2015 | Group3 |
| Y461 | Jack's Peak | Dhami MK, et al. | 2015 | Outgroup |
| Y466 | Soquel Valley | Dhami MK, et al. | 2015 | Group1 |
| Y467 | Soquel Valley | Dhami MK, et al. | 2015 | Group1 |
| Y514 | Soquel Valley | Dhami MK, et al. | 2015 | Group1 |
| Y515 | Soquel Valley | Dhami MK, et al. | 2015 | Group1 |
| Y520 | Soquel Valley | Dhami MK, et al. | 2015 | Group1 |
| Y556 | Jack's Peak | Dhami MK, et al. | 2015 | Group2 |
| Y557 | Jack's Peak | Dhami MK, et al. | 2015 | Group3 |
| Y577 | Jack's Peak | Dhami MK, et al. | 2015 | Group2 |
| Y620 | Jack's Peak | Dhami MK, et al. | 2015 | Group3 |
| Y633 | Jack's Peak | Dhami MK, et al. | 2015 | Group3 |
| Y641 | Jack's Peak | Dhami MK, et al. | 2015 | Group3 |
| Y642 | Jack's Peak | Dhami MK, et al. | 2015 | Group3 |
| Y643 | Jack's Peak | Dhami MK, et al. | 2015 | Group3 |
| Y644 | Jack's Peak | Dhami MK, et al. | 2015 | Group3 |
| Y680 | Carmel Highlands | Dhami MK, et al. | 2015 | Group3 |
| Y681 | Carmel Highlands | Dhami MK, et al. | 2015 | Group3 |
| Y75 | La Honda | Dhami MK, et al. | 2015 | Outgroup |
| Y76 | La Honda | Dhami MK, et al. | 2015 | Group3 |
| Y797 | Bodega Bay | Dhami MK, et al. | 2015 | Group3 |
| Y798 | Bodega Bay | Dhami MK, et al. | 2015 | Group3 |
| Y810 | Sweeney Ridge | Dhami MK, et al. | 2015 | Group3 |
| Y817 | Sweeney Ridge | Dhami MK, et al. | 2015 | Group2 |
| Y818 | Sweeney Ridge | Dhami MK, et al. | 2015 | Group2 |
| Y819 | Sweeney Ridge | Dhami MK, et al. | 2015 | Group3 |
| Y82 | La Honda | Dhami MK, et al. | 2015 | Group3 |
| Y820 | Sweeney Ridge | Dhami MK, et al. | 2015 | Group2 |
| Y821 | Sweeney Ridge | Dhami MK, et al. | 2015 | Group3 |
| Y825 | Sweeney Ridge | Dhami MK, et al. | 2015 | Group2 |
| Y837 | Sweeney Ridge | Dhami MK, et al. | 2015 | Group3 |
| Y843 | Sweeney Ridge | Dhami MK, et al. | 2015 | Group3 |
| Y844 | Sweeney Ridge | Dhami MK, et al. | 2015 | Group3 |
| Y845 | Sweeney Ridge | Dhami MK, et al. | 2015 | Group2 |
| Y846 | Sweeney Ridge | Dhami MK, et al. | 2015 | Group2 |
| Y850 | Sweeney Ridge | Dhami MK, et al. | 2015 | Group3 |
| Y851 | Sweeney Ridge | Dhami MK, et al. | 2015 | Group3 |
| Y856 | Sweeney Ridge | Dhami MK, et al. | 2015 | Group3 |
| Y857 | Sweeney Ridge | Dhami MK, et al. | 2015 | Group3 |
| Y858 | Sweeney Ridge | Dhami MK, et al. | 2015 | Group3 |
| Y859 | Sweeney Ridge | Dhami MK, et al. | 2015 | Group2 |
| Y864 | Sweeney Ridge | Dhami MK, et al. | 2015 | Group2 |
| Y865 | Sweeney Ridge | Dhami MK, et al. | 2015 | Group2 |
| Y866 | Sweeney Ridge | Dhami MK, et al. | 2015 | Group3 |
| Y867 | Sweeney Ridge | Dhami MK, et al. | 2015 | Group3 |
| Y883 | Sweeney Ridge | Dhami MK, et al. | 2015 | Group2 |
| Y884 | Sweeney Ridge | Dhami MK, et al. | 2015 | Group3 |
| Y885 | Sweeney Ridge | Dhami MK, et al. | 2015 | Group2 |
| Y886 | Sweeney Ridge | Dhami MK, et al. | 2015 | Group2 |
| Y893 | Sweeney Ridge | Dhami MK, et al. | 2015 | Group2 |
| Y894 | Sweeney Ridge | Dhami MK, et al. | 2015 | Group3 |
| Y895 | Sweeney Ridge | Dhami MK, et al. | 2015 | Group3 |
| Y896 | Sweeney Ridge | Dhami MK, et al. | 2015 | Group3 |
| Y914 | Swanton | Dhami MK, et al. | 2015 | Group3 |
| Y919 | Swanton | Dhami MK, et al. | 2015 | Group2 |
| Y920 | Swanton | Dhami MK, et al. | 2015 | Group2 |
| Y93 | La Honda | Dhami MK, et al. | 2015 | Group3 |
| Y939 | Swanton | Dhami MK, et al. | 2015 | Group3 |
| Y94 | La Honda | Dhami MK, et al. | 2015 | Group3 |
| Y940 | Swanton | Dhami MK, et al. | 2015 | Group2 |
| Y95 | La Honda | Dhami MK, et al. | 2015 | Group3 |
| Y96 | La Honda | Dhami MK, et al. | 2015 | Group3 |
| Y961 | Swanton | Dhami MK, et al. | 2015 | Group3 |
| Y968 | Swanton | Dhami MK, et al. | 2015 | Group3 |
| Y976 | Swanton | Dhami MK, et al. | 2015 | Group3 |
| Y982 | Swanton | Dhami MK, et al. | 2015 | Group3 |
| Y988 | Swanton | Dhami MK, et al. | 2015 | Group3 |
| Y989 | Swanton | Dhami MK, et al. | 2015 | Group3 |

#### **S2: Sequencing results**

| **Sample name** | **Raw reads** | **Filtered Reads** | **Total fragments** | **Mapped fragments** | **Assigned fragments** | **% Assigned fragments** |
| --- | --- | --- | --- | --- | --- | --- |
| fukami697 | 28190738 | 27766858 | 13883429 | 12886275 | 11474269 | 0.826472264 |
| fukami699 | 37803378 | 37198972 | 18599486 | 17302930 | 15716816 | 0.845013459 |
| fukami700 | 27894136 | 27693930 | 13846965 | 12907858 | 11715301 | 0.846055507 |
| fukami701 | 41757012 | 41343746 | 20671873 | 19222575 | 17565949 | 0.849751205 |
| fukami702 | 39398664 | 38773952 | 19386976 | 18018794 | 16213418 | 0.836304641 |
| fukami703 | 52725858 | 52029780 | 26014890 | 24158917 | 22097224 | 0.849406782 |
| fukami704 | 41828938 | 41407646 | 20703823 | 19267476 | 17632234 | 0.851641458 |
| fukami705 | 33322958 | 32383140 | 16191570 | 15083525 | 13821271 | 0.853609069 |
| fukami706 | 34109926 | 33778458 | 16889229 | 16549060 | 15069270 | 0.89224144 |
| fukami707 | 28286268 | 27904162 | 13952081 | 13722827 | 12227099 | 0.87636382 |
| fukami708 | 31943070 | 31368220 | 15684110 | 15433214 | 13781063 | 0.878664011 |
| fukami709 | 39662838 | 39359790 | 19679895 | 17003292 | 15912880 | 0.808585615 |
| fukami710 | 27144114 | 26211684 | 13105842 | 12895689 | 11585303 | 0.88398006 |
| fukami711 | 28166794 | 27640980 | 13820490 | 11800731 | 10867287 | 0.786317055 |
| fukami712 | 55309948 | 55048528 | 27524264 | 23929958 | 22345243 | 0.811837984 |
| fukami754 | 29792616 | 29548352 | 14774176 | 12700719 | 11747141 | 0.795113108 |
| fukami755 | 43468642 | 42738580 | 21369290 | 18821258 | 17408293 | 0.814640683 |
| fukami756 | 34724592 | 34409600 | 17204800 | 14989549 | 13868659 | 0.806092428 |
| fukami757 | 30832232 | 30506372 | 15253186 | 13178614 | 12220796 | 0.801196288 |
| fukami758 | 48039216 | 47341274 | 23670637 | 20127424 | 18512199 | 0.78207439 |
| fukami759 | 31756254 | 31481192 | 15740596 | 13726849 | 12759824 | 0.810631567 |
| fukami760 | 28528680 | 28367436 | 14183718 | 12325283 | 11492219 | 0.810240235 |
| fukami761 | 35682030 | 35115774 | 17557887 | 15366666 | 14229654 | 0.810442282 |
| fukami762 | 34283000 | 33430454 | 16715227 | 14609095 | 13440725 | 0.804100656 |
| fukami763 | 32364254 | 31861814 | 15930907 | 13501088 | 12404344 | 0.778633884 |
| fukami764 | 42500782 | 42072666 | 21036333 | 18148675 | 16726131 | 0.79510678 |
| fukami765 | 26608984 | 26142200 | 13071100 | 11296708 | 10450560 | 0.799516491 |
| fukami766 | 27211698 | 26668544 | 13334272 | 11619151 | 10756647 | 0.806691734 |
| fukami767 | 52732800 | 52002582 | 26001291 | 23002591 | 21279799 | 0.81841317 |
| fukami768 | 31611776 | 30808752 | 15404376 | 13362639 | 12351072 | 0.801789829 |
| fukami769 | 25248064 | 24733356 | 12366678 | 11943584 | 10819978 | 0.874930034 |
| fukami770 | 28095082 | 27785538 | 13892769 | 13688903 | 12382656 | 0.891302231 |
| fukami771 | 31408326 | 30950526 | 15475263 | 15306476 | 13445386 | 0.868830856 |
| fukami772 | 30744214 | 30520664 | 15260332 | 15157181 | 13633865 | 0.893418636 |
| fukami773 | 26130930 | 25640994 | 12820497 | 12729453 | 11401262 | 0.889299533 |
| fukami774 | 44943328 | 44529432 | 22264716 | 22110472 | 19754960 | 0.887276532 |
| fukami775 | 23725756 | 23480374 | 11740187 | 11612167 | 10235590 | 0.871842161 |
| fukami776 | 39898538 | 38975668 | 19487834 | 19292896 | 17034529 | 0.874110945 |
| fukami777 | 29094534 | 28791526 | 14395763 | 14081739 | 12614410 | 0.876258521 |
| fukami778 | 28331308 | 27471134 | 13735567 | 13408170 | 11925857 | 0.868246429 |
| fukami779 | 37375338 | 36862976 | 18431488 | 18040715 | 16174130 | 0.877527088 |
| fukami780 | 34029610 | 33060532 | 16530266 | 16233947 | 14619193 | 0.884389459 |
| fukami781 | 25626172 | 25321554 | 12660777 | 12392452 | 11144752 | 0.880258139 |
| fukami782 | 41238912 | 40312048 | 20156024 | 19746336 | 17755015 | 0.880878838 |
| fukami785 | 37159812 | 36559576 | 18279788 | 15818725 | 14374159 | 0.786341669 |
| fukami786 | 27317158 | 27026544 | 13513272 | 11459501 | 10522596 | 0.778686021 |
| fukami787 | 35460388 | 35121022 | 17560511 | 15219793 | 14046511 | 0.799891928 |
| fukami788 | 33696108 | 33300944 | 16650472 | 14237738 | 12983908 | 0.779792188 |
| fukami789 | 34793824 | 34247958 | 17123979 | 14592981 | 13384897 | 0.781646427 |
| fukami790 | 40089430 | 39477176 | 19738588 | 17347477 | 16094622 | 0.81538872 |
| fukami791 | 29631596 | 29170302 | 14585151 | 12464828 | 11413234 | 0.782524226 |
| fukami792 | 57630462 | 57329072 | 28664536 | 24972525 | 23176667 | 0.80854848 |
| fukami793 | 36048464 | 35637588 | 17818794 | 15590311 | 14417639 | 0.80912541 |
| fukami794 | 28042758 | 27828476 | 13914238 | 11939503 | 11048740 | 0.794060012 |
| fukami795 | 35289116 | 34836512 | 17418256 | 15323339 | 14347904 | 0.823727932 |
| fukami796 | 30906836 | 30656906 | 15328453 | 13476683 | 12497610 | 0.815321024 |
| fukami797 | 27220350 | 26776798 | 13388399 | 11405204 | 10487440 | 0.783322935 |
| fukami798 | 34859942 | 34473008 | 17236504 | 16973457 | 15338999 | 0.889913581 |
| fukami799 | 28486090 | 28247016 | 14123508 | 12276097 | 11442745 | 0.81019142 |
| fukami800 | 27130640 | 26157656 | 13078828 | 11096961 | 10177230 | 0.778145412 |
| fukami802 | 35507224 | 35175306 | 17587653 | 15650078 | 14605961 | 0.830466748 |
| fukami803 | 15487844 | 13943336 | 6971668 | 6839552 | 5993715 | 0.859724674 |
| fukami804 | 29572648 | 29071310 | 14535655 | 12517149 | 11619713 | 0.799393835 |
| fukami805 | 33628124 | 32919322 | 16459661 | 14497483 | 13587392 | 0.825496467 |
| fukami806 | 34843642 | 33791950 | 16895975 | 14685132 | 13633202 | 0.806890517 |
| fukami807 | 34243006 | 33396640 | 16698320 | 14811384 | 13793137 | 0.826019444 |
| fukami808 | 28791164 | 28518660 | 14259330 | 12467128 | 11625956 | 0.81532274 |
| fukami810 | 30007112 | 29304188 | 14652094 | 13436675 | 11938489 | 0.814797462 |
| fukami811 | 47067920 | 46685036 | 23342518 | 21577859 | 19579457 | 0.838789414 |
| fukami812 | 25048478 | 24269416 | 12134708 | 11162413 | 10052259 | 0.828389031 |
| fukami813 | 44733386 | 44162860 | 22081430 | 20551427 | 18637614 | 0.844040173 |
| fukami814 | 29607106 | 28202082 | 14101041 | 12979660 | 11624321 | 0.824359067 |
| fukami815 | 31593302 | 30338590 | 15169295 | 14150348 | 12600721 | 0.830672816 |
| fukami816 | 32283810 | 31734928 | 15867464 | 14559413 | 13046701 | 0.822229753 |
| fukami817 | 34460884 | 33906416 | 16953208 | 15633307 | 14321072 | 0.844741125 |
| fukami818 | 30993176 | 30609682 | 15304841 | 14170611 | 12843215 | 0.839160302 |
| fukami819 | 33632312 | 33414202 | 16707101 | 15554666 | 14389546 | 0.861283235 |
| fukami820 | 25932006 | 25517026 | 12758513 | 11765010 | 10752018 | 0.842732848 |
| fukami822 | 25636634 | 25250846 | 12625423 | 11820668 | 10897859 | 0.863167832 |
| fukami823 | 38727348 | 37924250 | 18962125 | 17418420 | 15877427 | 0.83732319 |
| fukami824 | 26415248 | 25853578 | 12926789 | 11942923 | 10952279 | 0.847254411 |
| fukami825 | 39886500 | 38757124 | 19378562 | 19082270 | 17372604 | 0.896485714 |
| fukami826 | 41141608 | 40506686 | 20253343 | 18756699 | 17129440 | 0.845758648 |
| fukami827 | 33189642 | 32690260 | 16345130 | 15277673 | 13737289 | 0.840451498 |
| fukami828 | 27014828 | 26642700 | 13321350 | 12425882 | 11175813 | 0.838939972 |
| fukami829 | 21756904 | 21480234 | 10740117 | 10200036 | 9285032 | 0.864518701 |
| fukami830 | 25651856 | 25427998 | 12713999 | 12006597 | 10850177 | 0.853403953 |
| fukami832 | 30940860 | 30623644 | 15311822 | 14368091 | 12925975 | 0.844182684 |
| fukami833 | 28310792 | 27824800 | 13912400 | 13680713 | 12361633 | 0.888533467 |
| fukami834 | 34225322 | 30591020 | 15295510 | 15019880 | 13692393 | 0.895190353 |
| fukami835 | 26136880 | 25913116 | 12956558 | 11204878 | 10315646 | 0.796171792 |
| fukami836 | 33073664 | 32368206 | 16184103 | 15970585 | 14594213 | 0.901762242 |
| fukami837 | 31591342 | 31157924 | 15578962 | 15334850 | 13937939 | 0.894664163 |
| fukami838 | 30254102 | 29857692 | 14928846 | 14692552 | 13340897 | 0.893632167 |
| fukami839 | 29667770 | 29205450 | 14602725 | 14389178 | 13137234 | 0.899642635 |
| fukami840 | 34559644 | 34203194 | 17101597 | 16832011 | 15352602 | 0.897729142 |
| fukami841 | 24048618 | 23364468 | 11682234 | 10945015 | 9909760 | 0.848276109 |
| fukami842 | 35735106 | 34960210 | 17480105 | 16068097 | 14342854 | 0.820524476 |
| fukami843 | 36787118 | 36316524 | 18158262 | 16717463 | 14949202 | 0.823272734 |
| fukami844 | 26972744 | 26421084 | 13210542 | 12189722 | 10920650 | 0.826661768 |
| fukami845 | 27014300 | 26345946 | 13172973 | 12166395 | 10908917 | 0.8281287 |
| fukami846 | 27089174 | 26677724 | 13338862 | 12354041 | 11158225 | 0.836520012 |
| fukami847 | 28747484 | 28521114 | 14260557 | 13201203 | 11949419 | 0.837934942 |
| fukami848 | 32243430 | 32090814 | 16045407 | 14909676 | 13603492 | 0.847812212 |
| fukami849 | 33123104 | 30059312 | 15029656 | 14723796 | 13493040 | 0.897761067 |
| fukami850 | 27689098 | 27300944 | 13650472 | 13354803 | 12075768 | 0.884641059 |
| fukami851 | 30887038 | 30603454 | 15301727 | 14971109 | 13565396 | 0.886527122 |
| fukami852 | 25431194 | 25074704 | 12537352 | 12286710 | 11172936 | 0.891171916 |
| fukami853 | 27908500 | 27605992 | 13802996 | 13508400 | 12305242 | 0.891490659 |
| fukami854 | 39121726 | 38902320 | 19451160 | 19096211 | 17458737 | 0.897567909 |
| fukami855 | 26287140 | 25910606 | 12955303 | 12669262 | 11541956 | 0.890905909 |
| fukami856 | 31136478 | 30424510 | 15212255 | 13167027 | 12182497 | 0.800834393 |
| fukami857 | 58496016 | 58225476 | 29112738 | 28722250 | 26219601 | 0.900622985 |
| fukami859 | 26061520 | 25483360 | 12741680 | 12568204 | 11314979 | 0.888028816 |
| fukami860 | 27306698 | 27008352 | 13504176 | 13289002 | 12040757 | 0.891632114 |
| fukami861 | 35990410 | 35677558 | 17838779 | 17543192 | 15788967 | 0.88509236 |
| fukami862 | 37240034 | 37074356 | 18537178 | 18237350 | 16444836 | 0.887127264 |
| fukami863 | 45401852 | 45196670 | 22598335 | 22293954 | 20379012 | 0.901792632 |
| fukami864 | 44639576 | 44326038 | 22163019 | 21850446 | 19891921 | 0.897527589 |
| fukami865 | 24781186 | 24429882 | 12214941 | 11801660 | 10547767 | 0.863513545 |
| fukami866 | 26000566 | 25595730 | 12797865 | 12309880 | 11022774 | 0.861297881 |
| fukami867 | 32493408 | 32049088 | 16024544 | 15453251 | 13743114 | 0.857629022 |
| fukami868 | 31593912 | 31278654 | 15639327 | 13391811 | 12447375 | 0.795902215 |
| fukami869 | 33854686 | 33052454 | 16526227 | 16260639 | 14608543 | 0.883961173 |
| fukami870 | 32593420 | 31269238 | 15634619 | 15096551 | 13688309 | 0.875512796 |
| fukami871 | 37148848 | 36692154 | 18346077 | 17780943 | 16093946 | 0.877241821 |
| fukami872 | 68491100 | 67575306 | 33787653 | 32996597 | 29921211 | 0.885566423 |
| fukami737 | 41200102 | 40499084 | 20249542 | 20047217 | 18213928 | 0.899473578 |
| fukami738 | 53987362 | 53328700 | 26664350 | 26365875 | 24024349 | 0.900991361 |
| fukami739 | 34260308 | 33763768 | 16881884 | 16626004 | 14957130 | 0.885987014 |
| fukami740 | 33769536 | 33307712 | 16653856 | 16489521 | 14960901 | 0.898344564 |
| fukami741 | 32459676 | 31957250 | 15978625 | 15789637 | 14401589 | 0.901303398 |
| fukami742 | 29384852 | 29194110 | 14597055 | 14435615 | 13240683 | 0.907079065 |
| fukami743 | 23275638 | 23060648 | 11530324 | 11421611 | 10451702 | 0.906453453 |
| fukami744 | 46658700 | 45997902 | 22998951 | 22774017 | 20722876 | 0.901035704 |
| fukami745 | 21126262 | 20666314 | 10333157 | 9948147 | 9028113 | 0.873703264 |
| fukami746 | 31844288 | 31602950 | 15801475 | 15179237 | 13783609 | 0.872298883 |
| fukami747 | 29495668 | 28768498 | 14384249 | 13824515 | 12415641 | 0.863141413 |
| fukami748 | 22248636 | 21785888 | 10892944 | 10456115 | 9341357 | 0.857560362 |
| fukami750 | 24642432 | 24280658 | 12140329 | 11652353 | 10471905 | 0.862571764 |
| fukami751 | 24603356 | 23979618 | 11989809 | 11498209 | 10291760 | 0.858375642 |
| fukami713 | 32431812 | 31475222 | 15737611 | 14605447 | 13345254 | 0.847984742 |
| fukami714 | 30970518 | 30415740 | 15207870 | 14159393 | 12939871 | 0.850866755 |
| fukami715 | 49882424 | 48623214 | 24311607 | 20879510 | 19371723 | 0.796809647 |
| fukami716 | 36645898 | 35707002 | 17853501 | 16544592 | 15023964 | 0.841513606 |
| fukami717 | 54137944 | 50536936 | 25268468 | 24906158 | 22618829 | 0.895140497 |
| fukami718 | 48326120 | 47069038 | 23534519 | 21803728 | 19880641 | 0.844743885 |
| fukami719 | 30068446 | 29683312 | 14841656 | 12844824 | 11849902 | 0.79842182 |
| fukami720 | 34833326 | 34380092 | 17190046 | 14796264 | 13681593 | 0.795902059 |
| fukami721 | 25721338 | 25455444 | 12727722 | 11667953 | 10668520 | 0.838211268 |
| fukami722 | 42486360 | 41501296 | 20750648 | 19358741 | 17780791 | 0.856878831 |
| fukami723 | 40833164 | 40487748 | 20243874 | 18792170 | 17052363 | 0.842346826 |
| fukami724 | 26124914 | 25669540 | 12834770 | 11926237 | 10761656 | 0.838476732 |
| fukami725 | 28001394 | 27389582 | 13694791 | 12608899 | 11493368 | 0.83925107 |
| fukami726 | 33509578 | 33058744 | 16529372 | 15494857 | 14101233 | 0.853101558 |
| fukami727 | 30714244 | 29953716 | 14976858 | 13717365 | 12438315 | 0.830502299 |
| fukami728 | 26479414 | 25733896 | 12866948 | 11781207 | 10671628 | 0.829383005 |
| fukami729 | 22688988 | 22377500 | 11188750 | 9428729 | 8672942 | 0.77514843 |
| fukami730 | 32803488 | 32393296 | 16196648 | 14013435 | 13029781 | 0.804473926 |
| fukami731 | 48654092 | 48194630 | 24097315 | 21028209 | 19444918 | 0.806932972 |
| fukami732 | 33311954 | 32492268 | 16246134 | 13551101 | 12325788 | 0.758690529 |
| fukami733 | 32384216 | 31793188 | 15896594 | 13779539 | 12714263 | 0.799810513 |
| fukami734 | 38446112 | 38032606 | 19016303 | 16124710 | 14811540 | 0.778886411 |
| fukami735 | 29675260 | 29418098 | 14709049 | 12789105 | 11747905 | 0.798685557 |
| fukami736 | 31804162 | 31246408 | 15623204 | 13476782 | 12321408 | 0.788660764 |
| fukami873 | 41361452 | 41130368 | 20565184 | 20282805 | 18171515 | 0.883605758 |
| fukami874 | 27719560 | 27367490 | 13683745 | 13481879 | 12258116 | 0.895815875 |
| fukami875 | 26633138 | 25508284 | 12754142 | 12242716 | 11008617 | 0.863140539 |
| fukami876 | 33208444 | 31128100 | 15564050 | 15318938 | 13318315 | 0.855710114 |
| fukami877 | 34635202 | 34471024 | 17235512 | 16990576 | 15159841 | 0.879570099 |
| fukami878 | 43968856 | 43746642 | 21873321 | 21547922 | 19267622 | 0.880873188 |
| fukami879 | 62113570 | 60295038 | 30147519 | 29506116 | 26696149 | 0.885517279 |
| fukami880 | 41786768 | 41274234 | 20637117 | 20319714 | 18043472 | 0.87432135 |
| fukami15 | 30369328 | 29535142 | 14767571 | 13644688 | 12358561 | 0.836871616 |
| fukami17 | 35566616 | 35271318 | 17635659 | 17396517 | 15823435 | 0.897240925 |
| fukami18 | 30259498 | 30068814 | 15034407 | 14842446 | 13599352 | 0.904548613 |
| fukami19 | 25424720 | 25299868 | 12649934 | 12479796 | 11411327 | 0.902085892 |
| fukami20 | 39774704 | 39391172 | 19695586 | 17134978 | 15958399 | 0.810252561 |
| fukami21 | 32559464 | 32297176 | 16148588 | 15924067 | 14585051 | 0.903178098 |
| fukami22 | 37695340 | 37421776 | 18710888 | 18420495 | 16693895 | 0.892202177 |
| fukami23 | 24057022 | 23926018 | 11963009 | 11794046 | 10809826 | 0.903604269 |
| fukami24 | 46594370 | 46060638 | 23030319 | 22705703 | 20716707 | 0.899540601 |
| fukami66 | 43544814 | 42394070 | 21197035 | 20885901 | 18173924 | 0.857380478 |
| fukami67 | 28231258 | 27940454 | 13970227 | 13751587 | 12294550 | 0.880053703 |
| fukami68 | 46994944 | 46315506 | 23157753 | 22804629 | 20396504 | 0.880763518 |
| fukami69 | 32514510 | 31906454 | 15953227 | 15741561 | 13973955 | 0.875932813 |
| fukami70 | 32071936 | 29740576 | 14870288 | 14648982 | 13125489 | 0.88266542 |
| fukami71 | 40121090 | 39799502 | 19899751 | 19622690 | 17702557 | 0.88958686 |
| fukami72 | 30800058 | 30394806 | 15197403 | 14919377 | 13310258 | 0.875824508 |
| fukami73 | 29466128 | 29042734 | 14521367 | 12870219 | 11701671 | 0.805824341 |
| fukami74 | 41405536 | 41002654 | 20501327 | 17581537 | 16200505 | 0.790217384 |
| fukami75 | 27537882 | 27078882 | 13539441 | 11992452 | 10869248 | 0.802784103 |
| fukami77 | 47869516 | 46916594 | 23458297 | 20731053 | 18937558 | 0.807286138 |
| fukami78 | 23625404 | 23083678 | 11541839 | 9951246 | 9169368 | 0.794446015 |
| fukami79 | 41797152 | 41522722 | 20761361 | 18369044 | 16954686 | 0.816646173 |
| fukami80 | 36928030 | 36606860 | 18303430 | 18043236 | 16331006 | 0.892237466 |
| fukami89 | 38401256 | 38045014 | 19022507 | 18628986 | 16845627 | 0.885562928 |
| fukami90 | 30708486 | 29765664 | 14882832 | 13213763 | 12201827 | 0.819859218 |
| fukami91 | 28420874 | 27956538 | 13978269 | 12040540 | 11135321 | 0.796616591 |
| fukami92 | 24181948 | 23849100 | 11924550 | 10318656 | 9481883 | 0.795156463 |
| fukami93 | 17820010 | 17019562 | 8509781 | 7537876 | 6872214 | 0.807566493 |
| fukami94 | 41966180 | 41531500 | 20765750 | 18368557 | 16878371 | 0.812798526 |
| fukami96 | 34831522 | 34662246 | 17331123 | 15245040 | 14172423 | 0.817744067 |
| fukami104 | 39584644 | 39020798 | 19510399 | 19202142 | 17426040 | 0.893166767 |
| fukami105 | 36041722 | 35709618 | 17854809 | 17627019 | 15954334 | 0.893559489 |
| fukami106 | 45559346 | 45070716 | 22535358 | 22210227 | 20011829 | 0.88801913 |
| fukami107 | 31788052 | 31132198 | 15566099 | 15347649 | 13846788 | 0.889547728 |
| fukami108 | 32191678 | 31652746 | 15826373 | 15586372 | 13919548 | 0.879515983 |
| fukami109 | 28348554 | 27456420 | 13728210 | 13521442 | 11633915 | 0.84744588 |
| fukami110 | 31877746 | 31265532 | 15632766 | 15427877 | 13744305 | 0.879198537 |
| fukami111 | 25018526 | 24405842 | 12202921 | 12056190 | 10816816 | 0.886412032 |
| fukami112 | 32493930 | 32141754 | 16070877 | 15823153 | 14451171 | 0.899214834 |
| fukami113 | 36176414 | 35214028 | 17607014 | 17077837 | 15162554 | 0.861165556 |
| fukami114 | 28728482 | 28503100 | 14251550 | 14117617 | 12493625 | 0.876650259 |
| fukami115 | 34491644 | 33950970 | 16975485 | 16447310 | 14411721 | 0.848972562 |
| fukami116 | 37665070 | 37148078 | 18574039 | 18276869 | 16184235 | 0.871336331 |
| fukami117 | 34137890 | 33655250 | 16827625 | 15696653 | 14236510 | 0.84602016 |
| fukami118 | 24314738 | 22414486 | 11207243 | 10729565 | 9498525 | 0.847534492 |
| fukami119 | 27242824 | 26863364 | 13431682 | 13233358 | 11954898 | 0.890052192 |
| fukami120 | 41030872 | 40545728 | 20272864 | 19635256 | 17529452 | 0.864675657 |
| fukami122 | 39854454 | 39059986 | 19529993 | 16784431 | 15395995 | 0.788325679 |
| fukami123 | 36052804 | 35526834 | 17763417 | 16290325 | 14937769 | 0.840928803 |
| fukami124 | 30455832 | 29981900 | 14990950 | 13144200 | 12011620 | 0.801258092 |
| fukami125 | 32648498 | 32120620 | 16060310 | 15030341 | 13581802 | 0.845674959 |
| fukami126 | 34485656 | 34043104 | 17021552 | 16769306 | 15244946 | 0.895626086 |
| fukami127 | 29632558 | 29271408 | 14635704 | 12825973 | 11726683 | 0.801238055 |
| fukami128 | 29544716 | 28932646 | 14466323 | 14228123 | 12973733 | 0.896823125 |
| fukami8 | 37705580 | 37226996 | 18613498 | 18130594 | 16386052 | 0.880331682 |
| fukami137 | 33459628 | 32701554 | 16350777 | 14451885 | 13403923 | 0.819772846 |
| fukami138 | 28236930 | 27471824 | 13735912 | 11981002 | 11071784 | 0.806046515 |
| fukami139 | 29000118 | 28405152 | 14202576 | 12178658 | 11142647 | 0.784551126 |
| fukami140 | 40745194 | 40126592 | 20063296 | 17548640 | 16108020 | 0.802860108 |
| fukami141 | 37586436 | 37130378 | 18565189 | 18135715 | 16569541 | 0.892505915 |
| fukami142 | 45299470 | 44824636 | 22412318 | 21930365 | 20050296 | 0.894610544 |
| fukami143 | 34271702 | 33897712 | 16948856 | 14776243 | 13501076 | 0.796577421 |
| fukami144 | 31263378 | 30607034 | 15303517 | 13426002 | 12157879 | 0.794449995 |
| fukami145 | 32321832 | 32070456 | 16035228 | 14232272 | 13372813 | 0.833964631 |
| fukami146 | 37840360 | 37544688 | 18772344 | 18385961 | 16836603 | 0.896883362 |
| fukami147 | 25863542 | 25734962 | 12867481 | 12577223 | 11517220 | 0.895064077 |
| fukami148 | 25217724 | 24962578 | 12481289 | 12259412 | 11129079 | 0.89166103 |
| fukami149 | 33537706 | 33060228 | 16530114 | 16139629 | 14682364 | 0.888219162 |
| fukami150 | 28029376 | 27568910 | 13784455 | 13522929 | 12083398 | 0.876595992 |
| fukami151 | 43313282 | 42909052 | 21454526 | 21005034 | 19063627 | 0.888559691 |
| fukami152 | 30778796 | 30561836 | 15280918 | 14890748 | 13654877 | 0.893590097 |
| fukami161 | 26784140 | 26532368 | 13266184 | 11772344 | 10919482 | 0.823106479 |
| fukami162 | 30724172 | 29831118 | 14915559 | 13017132 | 12053082 | 0.80808785 |
| fukami163 | 23320906 | 22983354 | 11491677 | 10110148 | 9327596 | 0.811682751 |
| fukami164 | 33799310 | 33335812 | 16667906 | 16415237 | 14890198 | 0.893345451 |
| fukami165 | 45068552 | 44583354 | 22291677 | 21531974 | 19289476 | 0.865321887 |
| fukami166 | 40643018 | 40067626 | 20033813 | 17092455 | 15835942 | 0.790460708 |
| fukami167 | 34799108 | 34313624 | 17156812 | 14562460 | 13430419 | 0.78280388 |
| fukami168 | 27127106 | 26488310 | 13244155 | 11613730 | 10732784 | 0.810378918 |
| fukami169 | 32411046 | 31959962 | 15979981 | 15745037 | 14235515 | 0.890834288 |
| fukami171 | 35617184 | 35229032 | 17614516 | 15341225 | 14167470 | 0.804306516 |
| fukami172 | 34159866 | 33931318 | 16965659 | 16682650 | 15006159 | 0.884501981 |
| fukami173 | 29857954 | 29496950 | 14748475 | 12801375 | 11918148 | 0.808093583 |
| fukami174 | 33509806 | 33100918 | 16550459 | 14399621 | 13428143 | 0.811345655 |
| fukami175 | 27547460 | 26969108 | 13484554 | 11762491 | 10583609 | 0.784869043 |
| fukami176 | 31317856 | 30791214 | 15395607 | 14173636 | 12924836 | 0.839514545 |
| fukami177 | 29801572 | 29646674 | 14823337 | 12798372 | 11769706 | 0.793998409 |
| fukami178 | 25933968 | 25670768 | 12835384 | 10933461 | 10126673 | 0.788965332 |
| fukami179 | 36631964 | 36420150 | 18210075 | 16015512 | 15094471 | 0.828907679 |
| fukami180 | 28354414 | 28017978 | 14008989 | 11890025 | 10945236 | 0.78130092 |
| fukami181 | 29448022 | 28926078 | 14463039 | 14253570 | 12862545 | 0.889339025 |
| fukami182 | 32480818 | 32076824 | 16038412 | 13947568 | 13056978 | 0.814106658 |
| fukami183 | 37307828 | 36993672 | 18496836 | 15952565 | 14895083 | 0.805277346 |
| fukami184 | 27824374 | 27581044 | 13790522 | 12708841 | 11610290 | 0.841903591 |
| fukami185 | 32592122 | 31727208 | 15863604 | 13940210 | 12947938 | 0.816204061 |
| fukami186 | 27759304 | 27331686 | 13665843 | 11602662 | 10667021 | 0.780560775 |
| fukami187 | 29392954 | 28866254 | 14433127 | 12128573 | 11109411 | 0.769716154 |
| fukami188 | 31345716 | 31065616 | 15532808 | 14324189 | 13015964 | 0.837965936 |
| fukami189 | 30487450 | 29899860 | 14949930 | 12989070 | 11907294 | 0.796478244 |
| fukami190 | 36269936 | 35967338 | 17983669 | 15611643 | 14443541 | 0.803147622 |
| fukami192 | 33786748 | 33623932 | 16811966 | 14263185 | 13206400 | 0.785535731 |
| fukami193 | 23769922 | 23369918 | 11684959 | 10196254 | 9434146 | 0.807375191 |
| fukami194 | 22828086 | 22011310 | 11005655 | 9638250 | 8921963 | 0.810670787 |
| fukami195 | 44813468 | 44414842 | 22207421 | 19554924 | 18429655 | 0.829887226 |
| fukami196 | 26593120 | 26248454 | 13124227 | 11501221 | 10633354 | 0.810208022 |
| fukami197 | 36445838 | 36014128 | 18007064 | 15329103 | 14234743 | 0.790508825 |
| fukami198 | 41609616 | 41308560 | 20654280 | 17892794 | 16698458 | 0.808474466 |
| fukami199 | 35226164 | 34082036 | 17041018 | 14849458 | 13761041 | 0.807524586 |
| fukami200 | 35407672 | 34077342 | 17038671 | 14728671 | 13591886 | 0.797708108 |
| fukami201 | 39128346 | 38845630 | 19422815 | 16619301 | 15328291 | 0.789189981 |
| fukami202 | 41013190 | 40338476 | 20169238 | 18873444 | 17192074 | 0.852390854 |
| fukami203 | 31590338 | 30198598 | 15099299 | 13964084 | 12759723 | 0.845053999 |
| fukami204 | 32289984 | 31931758 | 15965879 | 14770977 | 13552432 | 0.848837198 |
| fukami205 | 39712296 | 39426514 | 19713257 | 18276950 | 16798541 | 0.852144372 |
| fukami206 | 32465036 | 32041926 | 16020963 | 14833721 | 13583172 | 0.847837424 |
| fukami207 | 28096898 | 27767472 | 13883736 | 12926423 | 11910205 | 0.85785303 |
| fukami208 | 30786436 | 30233208 | 15116604 | 14060264 | 12825162 | 0.848415557 |
| fukami210 | 30321488 | 29748804 | 14874402 | 14615190 | 13299573 | 0.894124886 |
| fukami211 | 30870178 | 30126950 | 15063475 | 14810095 | 13329262 | 0.884872979 |
| fukami212 | 27161738 | 26845054 | 13422527 | 13127058 | 11750311 | 0.8754172 |
| fukami213 | 38798962 | 38509728 | 19254864 | 18894911 | 17119598 | 0.889105111 |
| fukami216 | 37648452 | 37039800 | 18519900 | 18139489 | 16283870 | 0.879263387 |
| fukami217 | 28679270 | 28319558 | 14159779 | 13268005 | 12092082 | 0.853973921 |
| fukami218 | 44211180 | 43495926 | 21747963 | 20303832 | 18413457 | 0.84667502 |
| fukami219 | 29694262 | 29349596 | 14674798 | 13550065 | 12257244 | 0.835258107 |
| fukami220 | 33824756 | 33477480 | 16738740 | 15486403 | 14070559 | 0.840598456 |
| fukami221 | 49630678 | 49240448 | 24620224 | 22731098 | 20705676 | 0.841002746 |
| fukami222 | 21560472 | 21366576 | 10683288 | 10052551 | 9223105 | 0.863320824 |
| fukami223 | 28495398 | 28077768 | 14038884 | 13184511 | 12013931 | 0.855761113 |
| fukami224 | 34805508 | 34162502 | 17081251 | 16032234 | 14396746 | 0.842839087 |
| fukami225 | 29024786 | 28549830 | 14274915 | 14099649 | 12731477 | 0.891877605 |
| fukami226 | 38880472 | 38373246 | 19186623 | 18966071 | 17313005 | 0.902347693 |
| fukami227 | 29064614 | 28574264 | 14287132 | 14113638 | 12554579 | 0.878733325 |
| fukami228 | 36627966 | 36096302 | 18048151 | 17650087 | 16092316 | 0.891632389 |
| fukami229 | 41728682 | 41150974 | 20575487 | 20334188 | 18511506 | 0.89968738 |
| fukami231 | 33267814 | 32975856 | 16487928 | 16244573 | 14695358 | 0.89127985 |
| fukami233 | 39318838 | 38706920 | 19353460 | 19035325 | 17053769 | 0.881174167 |
| fukami234 | 31641246 | 31288752 | 15644376 | 14422243 | 13116825 | 0.838437084 |
| fukami235 | 28676318 | 28127242 | 14063621 | 13722353 | 12506404 | 0.889273396 |
| fukami236 | 25173424 | 24675104 | 12337552 | 12065566 | 10781466 | 0.873874007 |
| fukami237 | 27354858 | 26526912 | 13263456 | 13067843 | 11213704 | 0.84545868 |
| fukami238 | 27227242 | 26375788 | 13187894 | 12999623 | 11353891 | 0.860932837 |
| fukami240 | 33739310 | 33077326 | 16538663 | 16257714 | 14689804 | 0.888209887 |
| fukami241 | 27239800 | 26664204 | 13332102 | 13113023 | 11831461 | 0.88744153 |
| fukami242 | 31990496 | 31736762 | 15868381 | 15664497 | 14224139 | 0.896382498 |
| fukami243 | 27263688 | 26815678 | 13407839 | 13241585 | 11944561 | 0.890863994 |
| fukami244 | 21789958 | 21538988 | 10769494 | 10634604 | 9661811 | 0.897146235 |
| fukami245 | 33897044 | 33097194 | 16548597 | 16311199 | 14832548 | 0.896302448 |
| fukami246 | 34107082 | 33617234 | 16808617 | 14590976 | 13510626 | 0.80379165 |
| fukami247 | 29662504 | 29001534 | 14500767 | 14243560 | 12778135 | 0.881204077 |
| fukami248 | 31171632 | 30562858 | 15281429 | 15048780 | 13548146 | 0.886575856 |
| fukami249 | 27034720 | 26513968 | 13256984 | 13002759 | 11644075 | 0.878335148 |
| fukami250 | 30473418 | 29751786 | 14875893 | 14630155 | 13175855 | 0.885718592 |
| fukami251 | 29395422 | 29077344 | 14538672 | 14268928 | 12778283 | 0.878916795 |
| fukami252 | 23097044 | 22567316 | 11283658 | 11069346 | 9855900 | 0.873466743 |
| fukami253 | 33549962 | 32846482 | 16423241 | 16101840 | 14312038 | 0.871450282 |
| fukami254 | 29550210 | 29001646 | 14500823 | 14210555 | 12613386 | 0.869839319 |
| fukami255 | 33219854 | 32597848 | 16298924 | 16072676 | 14375857 | 0.882012641 |
| fukami256 | 40064308 | 39436774 | 19718387 | 18948930 | 16747595 | 0.849338995 |
| fukami257 | 48098812 | 47556032 | 23778016 | 23430584 | 21316886 | 0.89649557 |
| fukami258 | 37630040 | 37184330 | 18592165 | 18312840 | 16640541 | 0.895029761 |
| fukami259 | 26193754 | 25807138 | 12903569 | 12643725 | 11347867 | 0.879436302 |
| fukami260 | 32931760 | 32287122 | 16143561 | 15917067 | 14638162 | 0.906749261 |
| fukami261 | 28939002 | 28436178 | 14218089 | 13977929 | 12710197 | 0.893945522 |
| fukami262 | 30907970 | 30151434 | 15075717 | 14828983 | 13416687 | 0.889953493 |
| fukami263 | 34968000 | 34524672 | 17262336 | 16907855 | 15079587 | 0.873554251 |
| fukami264 | 44970598 | 44536704 | 22268352 | 21968162 | 19963572 | 0.89649975 |
| fukami266 | 26547082 | 26309046 | 13154523 | 12983070 | 11715019 | 0.890569654 |
| fukami267 | 43343924 | 42677344 | 21338672 | 20993807 | 18744972 | 0.87845073 |
| fukami269 | 28638112 | 28038412 | 14019206 | 13732700 | 12099534 | 0.863068422 |
| fukami270 | 35590148 | 34763772 | 17381886 | 17136471 | 15272473 | 0.878643031 |
| fukami271 | 33040170 | 32119760 | 16059880 | 15822035 | 14063371 | 0.875683442 |
| fukami272 | 51383538 | 50879238 | 25439619 | 25107723 | 22799282 | 0.896211614 |
| fukami273 | 40500614 | 39759738 | 19879869 | 19524595 | 17597734 | 0.88520372 |
| fukami275 | 30622336 | 30459544 | 15229772 | 14032923 | 12689079 | 0.8331759 |
| fukami276 | 27640666 | 27258114 | 13629057 | 13322792 | 12126961 | 0.889787239 |
| fukami277 | 32034888 | 31699182 | 15849591 | 15664822 | 14447722 | 0.911551724 |
| fukami278 | 25939070 | 25556468 | 12778234 | 12610867 | 11482697 | 0.898613768 |
| fukami279 | 24986084 | 24633320 | 12316660 | 12114244 | 11021381 | 0.894835207 |
| fukami280 | 27828968 | 27278264 | 13639132 | 13387627 | 12058265 | 0.884093284 |
| fukami281 | 22576556 | 22309700 | 11154850 | 11026393 | 10055419 | 0.901439195 |
| fukami282 | 35391230 | 34768828 | 17384414 | 17081279 | 15371281 | 0.884198973 |
| fukami283 | 28834110 | 28529194 | 14264597 | 14087654 | 12590479 | 0.882638255 |
| fukami284 | 35757976 | 35403532 | 17701766 | 17439438 | 15669493 | 0.885193771 |
| fukami285 | 35042132 | 34382342 | 17191171 | 17004046 | 15541623 | 0.904046792 |
| fukami286 | 38383040 | 38113020 | 19056510 | 18855935 | 17338310 | 0.909836586 |
| fukami287 | 23852612 | 23391786 | 11695893 | 11485048 | 10215938 | 0.873463702 |
| fukami288 | 29574428 | 29076876 | 14538438 | 14351984 | 13031811 | 0.896369404 |
| fukami297 | 31115006 | 30473792 | 15236896 | 2351208 | 2216004 | 0.145436708 |
| fukami298 | 26958404 | 26272658 | 13136329 | 2057962 | 1852717 | 0.141037652 |
| fukami299 | 40232776 | 39283866 | 19641933 | 3006767 | 2751876 | 0.140102097 |
| fukami300 | 34163422 | 33704182 | 16852091 | 2583362 | 2444304 | 0.145044553 |
| fukami301 | 27156260 | 26706076 | 13353038 | 2148120 | 2028292 | 0.151897418 |
| fukami302 | 38903578 | 37916918 | 18958459 | 3005062 | 2847819 | 0.150213633 |
| fukami303 | 31186422 | 30778300 | 15389150 | 2380067 | 2246197 | 0.145959783 |
| fukami304 | 31209348 | 30878194 | 15439097 | 2490571 | 2333071 | 0.151114473 |
| fukami305 | 35684276 | 35039476 | 17519738 | 16365693 | 14970290 | 0.854481386 |
| fukami306 | 26591346 | 26065274 | 13032637 | 12219926 | 11067096 | 0.84918317 |
| fukami308 | 33171756 | 32987042 | 16493521 | 15419640 | 14207267 | 0.861384722 |
| fukami310 | 21583782 | 20657168 | 10328584 | 9673292 | 8726251 | 0.844864214 |
| fukami311 | 23732142 | 21584324 | 10792162 | 9957536 | 9120743 | 0.845126584 |
| fukami312 | 36116676 | 35883208 | 17941604 | 16769453 | 15440773 | 0.860612741 |
| fukami313 | 28900276 | 28521714 | 14260857 | 13484653 | 12380520 | 0.868146984 |
| fukami314 | 26355706 | 25653526 | 12826763 | 12133434 | 11105704 | 0.865822811 |
| fukami315 | 29541498 | 29054328 | 14527164 | 13575279 | 12466797 | 0.858171423 |
| fukami316 | 32433256 | 31516694 | 15758347 | 14892648 | 13624107 | 0.864564475 |
| fukami317 | 31379232 | 30659002 | 15329501 | 14308735 | 13118031 | 0.855737639 |
| fukami319 | 23299678 | 22785336 | 11392668 | 10589357 | 9730415 | 0.854094493 |
| fukami320 | 27569498 | 27146268 | 13573134 | 12740393 | 11819249 | 0.870782606 |
| fukami322 | 28761162 | 28241112 | 14120556 | 13681815 | 12341554 | 0.874013318 |
| fukami323 | 26444198 | 26057426 | 13028713 | 12040261 | 11074542 | 0.850010435 |
| fukami324 | 44190962 | 43988852 | 21994426 | 21719518 | 19924649 | 0.905895385 |
| fukami325 | 32440052 | 32205074 | 16102537 | 15980173 | 14199888 | 0.881841663 |
| fukami326 | 35577102 | 34224384 | 17112192 | 16866978 | 15301064 | 0.894161543 |
| fukami327 | 31838556 | 31403564 | 15701782 | 15461540 | 14030788 | 0.89357934 |
| fukami328 | 38034482 | 37480506 | 18740253 | 15869427 | 14652767 | 0.781887363 |
| fukami329 | 28496406 | 27908558 | 13954279 | 12956920 | 11865221 | 0.850292659 |
| fukami331 | 34404888 | 34172456 | 17086228 | 16022911 | 14763555 | 0.864061688 |
| fukami335 | 28098686 | 27378198 | 13689099 | 12632495 | 11436707 | 0.835460902 |
| fukami336 | 27192916 | 26450756 | 13225378 | 11556680 | 10643955 | 0.804812913 |
| fukami337 | 25080374 | 24590350 | 12295175 | 11312344 | 10323645 | 0.839650107 |
| fukami339 | 38526982 | 38049566 | 19024783 | 17632335 | 16122704 | 0.847457971 |
| fukami340 | 32153894 | 31332704 | 15666352 | 15420505 | 14075555 | 0.898457726 |
| fukami341 | 40985338 | 40399026 | 20199513 | 18686107 | 17135070 | 0.848291243 |
| fukami342 | 31136036 | 30553142 | 15276571 | 14200251 | 13063253 | 0.855116832 |
| fukami343 | 27914396 | 27497928 | 13748964 | 11600467 | 10719808 | 0.779681145 |
| fukami344 | 22950288 | 22672234 | 11336117 | 11158454 | 10095233 | 0.890537121 |
| fukami345 | 30267968 | 29903166 | 14951583 | 14754995 | 13360214 | 0.893565183 |
| fukami346 | 36736178 | 36150984 | 18075492 | 17905011 | 16290912 | 0.901270737 |
| fukami347 | 33514744 | 32476524 | 16238262 | 16034240 | 14560754 | 0.896694117 |
| fukami348 | 39999108 | 39800298 | 19900149 | 19689489 | 17920038 | 0.90049768 |
| fukami349 | 39938472 | 39281710 | 19640855 | 19407978 | 17572795 | 0.894706213 |
| fukami351 | 30118800 | 29856370 | 14928185 | 14694790 | 13341070 | 0.893683325 |
| fukami352 | 39634280 | 39193216 | 19596608 | 19393985 | 17755482 | 0.906048741 |
| fukami353 | 26159662 | 25578544 | 12789272 | 12570806 | 11195304 | 0.875366792 |
| fukami354 | 36879026 | 34838768 | 17419384 | 17153546 | 15443609 | 0.88657607 |
| fukami355 | 34149362 | 33754146 | 16877073 | 15627006 | 14378628 | 0.851962186 |
| fukami357 | 32526566 | 30908476 | 15454238 | 15196422 | 13652923 | 0.883442005 |
| fukami359 | 35400164 | 34822486 | 17411243 | 17054253 | 15028196 | 0.863131713 |
| fukami361 | 30610682 | 30112902 | 15056451 | 13960894 | 12795201 | 0.849815205 |
| fukami362 | 26213392 | 25518968 | 12759484 | 11847993 | 10857538 | 0.850938643 |
| fukami363 | 23883996 | 23419442 | 11709721 | 10750474 | 9714511 | 0.829610799 |
| fukami364 | 28859390 | 28454406 | 14227203 | 13040173 | 11759300 | 0.826536319 |
| fukami365 | 32328484 | 31922192 | 15961096 | 13654854 | 12708347 | 0.796207666 |
| fukami366 | 44310436 | 44113128 | 22056564 | 20523036 | 18921487 | 0.85786195 |
| fukami367 | 37190088 | 35862872 | 17931436 | 17316265 | 15661548 | 0.873412927 |
| fukami369 | 43309978 | 42910320 | 21455160 | 20405262 | 18613484 | 0.867552794 |
| fukami370 | 33305936 | 32987784 | 16493892 | 15689363 | 14279124 | 0.865721929 |
| fukami371 | 25419652 | 24683102 | 12341551 | 11696734 | 10532404 | 0.853410078 |
| fukami372 | 37184660 | 36523308 | 18261654 | 17300746 | 15677020 | 0.858466599 |
| fukami374 | 58800440 | 58157592 | 29078796 | 26784413 | 24213920 | 0.832700226 |
| fukami375 | 27622772 | 27133288 | 13566644 | 12844327 | 11658143 | 0.859324016 |
| fukami376 | 23447352 | 22950206 | 11475103 | 10901308 | 9817672 | 0.855562865 |
| fukami377 | 33538214 | 32849744 | 16424872 | 16023334 | 14375316 | 0.875216318 |
| fukami378 | 46705246 | 46515644 | 23257822 | 21726468 | 20106190 | 0.864491525 |
| fukami379 | 35902136 | 35775430 | 17887715 | 16698296 | 15383456 | 0.86000118 |
| fukami380 | 33482834 | 33029946 | 16514973 | 16272634 | 14814301 | 0.897022417 |
| fukami381 | 24996270 | 24315266 | 12157633 | 11333717 | 10390894 | 0.854680677 |
| fukami383 | 31017360 | 30475448 | 15237724 | 14151193 | 13026009 | 0.854852667 |
| fukami384 | 27387572 | 26864236 | 13432118 | 12521882 | 11514285 | 0.857220358 |
| fukami385 | 53095646 | 52801514 | 26400757 | 24617827 | 22525308 | 0.853206899 |
| fukami386 | 45514998 | 44936408 | 22468204 | 20840987 | 18930293 | 0.842536991 |
| fukami388 | 30743492 | 30312220 | 15156110 | 13887575 | 12401650 | 0.818260754 |
| fukami389 | 33578308 | 32933730 | 16466865 | 15227979 | 13973877 | 0.848605791 |
| fukami390 | 33375990 | 32991130 | 16495565 | 14254035 | 13125452 | 0.795695813 |
| fukami391 | 25690018 | 25328708 | 12664354 | 11709557 | 10487407 | 0.828104379 |
| fukami392 | 34034642 | 33865252 | 16932626 | 15661823 | 14214709 | 0.839486386 |
| fukami393 | 39125330 | 38761188 | 19380594 | 19172354 | 17675370 | 0.912013842 |
| fukami394 | 34444102 | 33684282 | 16842141 | 16610005 | 15126187 | 0.898115447 |
| fukami395 | 16365060 | 16121226 | 8060613 | 7916644 | 7156302 | 0.887811138 |
| fukami396 | 28831906 | 28329500 | 14164750 | 13993965 | 12858413 | 0.907775499 |
| fukami397 | 29344730 | 28999808 | 14499904 | 14306080 | 13055586 | 0.900391203 |
| fukami398 | 28971074 | 28225574 | 14112787 | 13940700 | 12714631 | 0.900929845 |
| fukami399 | 34551778 | 34348030 | 17174015 | 16946132 | 15451028 | 0.89967477 |
| fukami400 | 38083560 | 37674652 | 18837326 | 18615809 | 17085652 | 0.907010475 |
| fukami401 | 33094054 | 32498320 | 16249160 | 15986834 | 14465728 | 0.890244665 |
| fukami402 | 42855246 | 42322308 | 21161154 | 20833177 | 18936086 | 0.894851292 |
| fukami403 | 30421410 | 29983204 | 14991602 | 14821129 | 13579034 | 0.905776047 |
| fukami404 | 27139214 | 26973278 | 13486639 | 13328216 | 12202426 | 0.90477887 |
| fukami405 | 26965614 | 26759092 | 13379546 | 13218487 | 12097263 | 0.904160948 |
| fukami406 | 26811162 | 26668232 | 13334116 | 13168811 | 12008755 | 0.90060376 |
| fukami407 | 29272698 | 28971890 | 14485945 | 14302964 | 12996314 | 0.897167151 |
| fukami408 | 43884298 | 41995326 | 20997663 | 18067264 | 16530101 | 0.787235275 |
| fukami409 | 29267686 | 29125976 | 14562988 | 13577853 | 12501452 | 0.85844004 |
| fukami413 | 42161362 | 41921754 | 20960877 | 19398897 | 17862653 | 0.852190154 |
| fukami414 | 32642596 | 31983804 | 15991902 | 14926652 | 13336869 | 0.833976409 |
| fukami415 | 24782966 | 24197754 | 12098877 | 10545952 | 9199174 | 0.76033288 |
| fukami416 | 34069582 | 33385116 | 16692558 | 15187541 | 13444289 | 0.805406158 |
| fukami417 | 34518544 | 34143748 | 17071874 | 15774057 | 14369871 | 0.841727803 |
| fukami419 | 40973394 | 40758638 | 20379319 | 18879076 | 17290090 | 0.848413531 |
| fukami420 | 35801394 | 34983556 | 17491778 | 16260849 | 14856476 | 0.84934053 |
| fukami421 | 29281292 | 27885304 | 13942652 | 13722811 | 12265061 | 0.879679203 |
| fukami422 | 24312862 | 23809742 | 11904871 | 11122262 | 10021625 | 0.841808786 |
| fukami423 | 41377332 | 41074322 | 20537161 | 19044757 | 17463747 | 0.850348644 |
| fukami424 | 33353854 | 32144948 | 16072474 | 13832179 | 12783082 | 0.795340033 |
| fukami425 | 31723172 | 31283842 | 15641921 | 15430326 | 14026851 | 0.896747337 |
| fukami426 | 42297410 | 42076228 | 21038114 | 20764488 | 19029694 | 0.904534218 |
| fukami427 | 34920714 | 34611120 | 17305560 | 17040384 | 15444501 | 0.892458898 |
| fukami428 | 29088690 | 28417508 | 14208754 | 14028611 | 12806766 | 0.901329279 |
| fukami429 | 36711090 | 35861282 | 17930641 | 17681269 | 16061609 | 0.895763236 |
| fukami430 | 34062902 | 33748934 | 16874467 | 16620105 | 15086210 | 0.894025868 |
| fukami431 | 32787542 | 32449620 | 16224810 | 15932083 | 14094698 | 0.868712669 |
| fukami432 | 27487244 | 25999784 | 12999892 | 12137071 | 10921241 | 0.840102441 |
| fukami433 | 32931708 | 32669794 | 16334897 | 16024757 | 14513725 | 0.888510347 |
| fukami434 | 30794318 | 30440406 | 15220203 | 14977444 | 13531287 | 0.889034594 |
| fukami435 | 46446086 | 45953906 | 22976953 | 22590786 | 20396914 | 0.887711874 |
| fukami436 | 40030898 | 39676910 | 19838455 | 19514609 | 17743330 | 0.894390717 |
| fukami437 | 32053826 | 31625530 | 15812765 | 15560949 | 14128148 | 0.893464742 |
| fukami438 | 23111566 | 22976594 | 11488297 | 11307591 | 10301977 | 0.896736653 |
| fukami439 | 39819642 | 39375756 | 19687878 | 19391279 | 17563032 | 0.892073386 |
| fukami440 | 47107296 | 46509572 | 23254786 | 22861353 | 20702411 | 0.890243024 |
| fukami441 | 39408268 | 38617382 | 19308691 | 18981262 | 17182859 | 0.889902842 |
| fukami442 | 34297772 | 33860708 | 16930354 | 16673354 | 15175724 | 0.896361884 |
| fukami443 | 34238392 | 33332090 | 16666045 | 16443122 | 14937140 | 0.896261831 |
| fukami444 | 38982876 | 38461360 | 19230680 | 18964053 | 17363685 | 0.90291581 |
| fukami445 | 42275804 | 41591614 | 20795807 | 20476234 | 18634578 | 0.896073809 |
| fukami446 | 26543844 | 25950630 | 12975315 | 12767374 | 11695077 | 0.9013328 |
| fukami447 | 52758682 | 52463424 | 26231712 | 25862646 | 23604343 | 0.899839972 |
| fukami448 | 28661318 | 28308144 | 14154072 | 13897237 | 12070364 | 0.852783849 |
| fukami450 | 28463616 | 28096226 | 14048113 | 13808454 | 12542050 | 0.892792505 |
| fukami451 | 39552454 | 39139896 | 19569948 | 19243765 | 17517661 | 0.895130687 |
| fukami452 | 41104266 | 40697628 | 20348814 | 20052444 | 18385354 | 0.903509856 |
| fukami453 | 31967284 | 31721926 | 15860963 | 15617970 | 14315129 | 0.902538452 |
| fukami454 | 27693010 | 27330134 | 13665067 | 13424758 | 12188546 | 0.891949231 |
| fukami455 | 34942112 | 34722784 | 17361392 | 17097322 | 15706291 | 0.904667725 |
| fukami456 | 36193184 | 36029982 | 18014991 | 17720547 | 16192928 | 0.898858512 |
| fukami457 | 36250994 | 34796272 | 17398136 | 17104576 | 15111385 | 0.868563448 |
| fukami458 | 32983376 | 32215462 | 16107731 | 15809615 | 13846863 | 0.859640815 |
| fukami459 | 31770780 | 30711636 | 15355818 | 15057452 | 12678949 | 0.825677212 |
| fukami460 | 45935402 | 45554218 | 22777109 | 22307556 | 20242821 | 0.888735309 |
| fukami461 | 31212314 | 30100908 | 15050454 | 14754861 | 12357179 | 0.821050249 |
| fukami462 | 28180342 | 27375594 | 13687797 | 12676794 | 11625820 | 0.849356547 |
| fukami463 | 30433062 | 29487616 | 14743808 | 14467168 | 12586403 | 0.853673827 |
| fukami464 | 31297014 | 30827206 | 15413603 | 14266264 | 12975955 | 0.841850864 |
| fukami466 | 35678226 | 35005610 | 17502805 | 16219764 | 14538595 | 0.830643717 |
| fukami467 | 32892858 | 32623102 | 16311551 | 14998014 | 13867437 | 0.85016054 |
| fukami468 | 30169140 | 29753018 | 14876509 | 13614375 | 12480590 | 0.838946153 |
| fukami469 | 53557452 | 53306332 | 26653166 | 24654783 | 22846246 | 0.857168188 |
| fukami470 | 30971182 | 30074898 | 15037449 | 13906128 | 12263940 | 0.815559873 |
| fukami471 | 33106122 | 32627156 | 16313578 | 14969270 | 13769769 | 0.84406799 |
| fukami472 | 34360304 | 33538972 | 16769486 | 15521810 | 13883702 | 0.827914582 |
| fukami473 | 49071792 | 48654276 | 24327138 | 22511249 | 20613307 | 0.84733794 |
| fukami475 | 38280926 | 37787230 | 18893615 | 17330640 | 15991301 | 0.846386517 |
| fukami476 | 29024018 | 28057132 | 14028566 | 13010285 | 11336444 | 0.808097136 |
| fukami477 | 32373770 | 32027240 | 16013620 | 14845265 | 13676158 | 0.85403288 |
| fukami478 | 30617384 | 30295978 | 15147989 | 13908344 | 12802006 | 0.84512908 |
| fukami479 | 53891558 | 52806228 | 26403114 | 24427002 | 22390836 | 0.848037697 |
| fukami480 | 25907334 | 25243918 | 12621959 | 11679318 | 10446778 | 0.827666926 |
| fukami481 | 69363972 | 66389988 | 33194994 | 32661268 | 29479525 | 0.888071406 |
| fukami482 | 38611866 | 37918846 | 18959423 | 17530521 | 15647318 | 0.825305601 |
| fukami483 | 27138106 | 26189420 | 13094710 | 12876130 | 11655083 | 0.890060414 |
| fukami484 | 31921000 | 31062552 | 15531276 | 14858892 | 13361525 | 0.860297956 |
| fukami485 | 30855388 | 30539210 | 15269605 | 14101301 | 13149328 | 0.861143952 |
| fukami486 | 33623096 | 33026648 | 16513324 | 15306005 | 13746233 | 0.832432828 |
| fukami487 | 45621966 | 45213070 | 22606535 | 20863022 | 19373708 | 0.856995908 |
| fukami488 | 34055002 | 33641108 | 16820554 | 15441501 | 14222768 | 0.845558832 |
| fukami489 | 31694846 | 31351868 | 15675934 | 14983442 | 13558137 | 0.864901383 |
| fukami490 | 36057174 | 35795832 | 17897916 | 17175464 | 15420461 | 0.861578577 |
| fukami491 | 33275564 | 33093302 | 16546651 | 15870196 | 14267242 | 0.862243484 |
| fukami492 | 31505904 | 31299592 | 15649796 | 15067327 | 13604556 | 0.869312034 |
| fukami493 | 22951052 | 22756474 | 11378237 | 10841695 | 9818175 | 0.86289071 |
| fukami494 | 48920660 | 48401718 | 24200859 | 23098063 | 20823624 | 0.860449788 |
| fukami495 | 32197804 | 31262698 | 15631349 | 14415994 | 13224713 | 0.846037856 |
| fukami496 | 48252684 | 47385844 | 23692922 | 21905507 | 20003793 | 0.844294047 |
| fukami497 | 31087196 | 30619208 | 15309604 | 14210806 | 12901456 | 0.842703443 |
| fukami498 | 33071684 | 31940114 | 15970057 | 15423320 | 13893515 | 0.869972787 |
| fukami499 | 32225320 | 31243786 | 15621893 | 15307748 | 13430363 | 0.859714184 |
| fukami500 | 22511796 | 21908788 | 10954394 | 10732350 | 9336801 | 0.852333867 |
| fukami501 | 32998424 | 32211924 | 16105962 | 15797978 | 13817120 | 0.857888526 |
| fukami502 | 31439016 | 30603242 | 15301621 | 14207248 | 12941558 | 0.845763857 |
| fukami503 | 36643216 | 36152240 | 18076120 | 17712055 | 16101779 | 0.890776284 |
| fukami504 | 30884834 | 30353040 | 15176520 | 14876363 | 13141110 | 0.865884274 |
| fukami505 | 63103248 | 61423272 | 30711636 | 28466764 | 25980259 | 0.845941877 |
| fukami506 | 34400088 | 33848696 | 16924348 | 14498869 | 13380572 | 0.790610782 |
| fukami507 | 27710718 | 27093352 | 13546676 | 12686902 | 11379055 | 0.839988718 |
| fukami508 | 23566286 | 22993208 | 11496604 | 10738156 | 9484791 | 0.825008063 |
| fukami509 | 28624032 | 28082006 | 14041003 | 13798455 | 12489928 | 0.889532464 |
| fukami510 | 21297176 | 20717284 | 10358642 | 9666730 | 8341216 | 0.805242232 |
| fukami511 | 51476278 | 50483256 | 25241628 | 24716459 | 22453070 | 0.88952543 |
| fukami512 | 30551762 | 30238386 | 15119193 | 14012745 | 12838276 | 0.849137649 |
| fukami513 | 36810230 | 36494092 | 18247046 | 17926193 | 16422112 | 0.899987428 |
| fukami514 | 36355854 | 35590038 | 17795019 | 16587350 | 15201764 | 0.854270737 |
| fukami515 | 29896614 | 28974852 | 14487426 | 12531511 | 11619563 | 0.802044683 |
| fukami516 | 41762920 | 41485526 | 20742763 | 20394282 | 18284008 | 0.881464441 |
| fukami517 | 26421950 | 25689404 | 12844702 | 12631065 | 11464348 | 0.892535148 |
| fukami518 | 29413670 | 27846682 | 13923341 | 11994866 | 11075771 | 0.795482277 |
| fukami519 | 30072330 | 29839670 | 14919835 | 14683576 | 13203033 | 0.884931569 |
| fukami520 | 33698736 | 33034860 | 16517430 | 16246113 | 14545346 | 0.880605881 |
| fukami521 | 28320410 | 27768884 | 13884442 | 12922294 | 11363679 | 0.818446935 |
| fukami522 | 43145992 | 42449800 | 21224900 | 18214445 | 16784429 | 0.790789544 |
| fukami523 | 39089614 | 38457434 | 19228717 | 17953006 | 16025557 | 0.833417903 |
| fukami524 | 42836694 | 42479204 | 21239602 | 19788047 | 18220647 | 0.857861979 |
| fukami525 | 32745244 | 31917772 | 15958886 | 14898309 | 13411539 | 0.840380651 |
| fukami526 | 42176550 | 41001718 | 20500859 | 20088822 | 18283445 | 0.891837996 |
| fukami527 | 35927770 | 35436596 | 17718298 | 17334059 | 15647078 | 0.883102768 |
| fukami528 | 37162016 | 36760514 | 18380257 | 16935527 | 15437130 | 0.839875634 |
| fukami530 | 35724700 | 35540198 | 17770099 | 16242664 | 14792795 | 0.832454282 |
| fukami532 | 44185204 | 43375034 | 21687517 | 19948496 | 18057992 | 0.832644512 |
| fukami533 | 26410730 | 26256684 | 13128342 | 12214602 | 11234042 | 0.855709122 |
| fukami534 | 21590572 | 20970446 | 10485223 | 9052461 | 8268057 | 0.788543744 |
| fukami535 | 29236170 | 28922066 | 14461033 | 13387396 | 12172943 | 0.84177548 |
| fukami536 | 58622168 | 57197726 | 28598863 | 28084459 | 25356598 | 0.886629584 |
| fukami537 | 38231994 | 38013492 | 19006746 | 17623761 | 16115284 | 0.847871803 |
| fukami538 | 31724856 | 31186304 | 15593152 | 13421612 | 12388633 | 0.794491903 |
| fukami540 | 24583218 | 23833978 | 11916989 | 11109099 | 9721194 | 0.815742466 |
| fukami541 | 33195042 | 32936728 | 16468364 | 15384342 | 14119411 | 0.857365735 |
| fukami543 | 35870528 | 35300742 | 17650371 | 16317203 | 14956056 | 0.847350801 |
| fukami544 | 41526210 | 41240494 | 20620247 | 19256091 | 17704230 | 0.858584769 |
| fukami545 | 43066030 | 42794144 | 21397072 | 20946356 | 18767921 | 0.877125665 |
| fukami546 | 33547798 | 33000904 | 16500452 | 16187499 | 14500676 | 0.878804775 |
| fukami547 | 34411738 | 34179744 | 17089872 | 16746325 | 15067534 | 0.881664532 |
| fukami548 | 42267132 | 40312442 | 20156221 | 19847028 | 17801285 | 0.883165798 |
| fukami549 | 49363958 | 47924854 | 23962427 | 22221873 | 20327388 | 0.848302553 |
| fukami550 | 29745664 | 29543544 | 14771772 | 14493437 | 13163590 | 0.891131409 |
| fukami551 | 35954566 | 35345158 | 17672579 | 17273352 | 15461877 | 0.874907788 |
| fukami552 | 34037024 | 33483788 | 16741894 | 16420476 | 14910678 | 0.890620739 |
| fukami553 | 33904484 | 32688014 | 16344007 | 15077548 | 13686860 | 0.837423773 |
| fukami554 | 32395086 | 31601368 | 15800684 | 15301853 | 13795592 | 0.873100937 |
| fukami555 | 35113568 | 34799580 | 17399790 | 16843505 | 15354042 | 0.882426857 |
| fukami556 | 32630692 | 31387140 | 15693570 | 15417643 | 13829481 | 0.88121957 |
| fukami557 | 47632460 | 47435466 | 23717733 | 23026800 | 21151264 | 0.891791134 |
| fukami558 | 32258746 | 31850140 | 15925070 | 15413965 | 14086298 | 0.884536018 |
| fukami559 | 33279740 | 33009006 | 16504503 | 15930408 | 14499692 | 0.878529453 |
| fukami560 | 32653590 | 32426534 | 16213267 | 15676113 | 14305998 | 0.882363684 |
| fukami561 | 37487202 | 37044276 | 18522138 | 17237464 | 15841431 | 0.85527011 |
| fukami562 | 33019934 | 32628516 | 16314258 | 15145556 | 13907884 | 0.852498716 |
| fukami564 | 41827598 | 41359874 | 20679937 | 19232012 | 17644765 | 0.853231081 |
| fukami565 | 36037388 | 35831082 | 17915541 | 16633986 | 15223644 | 0.849745146 |
| fukami566 | 45950206 | 45724578 | 22862289 | 21254178 | 19513553 | 0.853525778 |
| fukami567 | 32283692 | 31841116 | 15920558 | 15606472 | 14096490 | 0.88542688 |
| fukami568 | 28457638 | 28053050 | 14026525 | 13430857 | 12025855 | 0.857365242 |
| fukami569 | 44101136 | 43754158 | 21877079 | 21436817 | 19366266 | 0.885230885 |
| fukami570 | 51610900 | 51291140 | 25645570 | 25136866 | 22956229 | 0.895134286 |
| fukami571 | 36407098 | 36031850 | 18015925 | 17670070 | 16101460 | 0.893734848 |
| fukami573 | 40741658 | 40182380 | 20091190 | 19664074 | 17863493 | 0.889120704 |
| fukami574 | 45330588 | 44815370 | 22407685 | 21921071 | 19973102 | 0.891350534 |
| fukami575 | 47579486 | 47302188 | 23651094 | 23150200 | 20989904 | 0.887481315 |
| fukami576 | 46083378 | 45832018 | 22916009 | 22431082 | 20449845 | 0.892382482 |
| fukami577 | 56189876 | 55846752 | 27923376 | 26020166 | 24098740 | 0.863031032 |
| fukami578 | 47049056 | 46678530 | 23339265 | 21734971 | 20112955 | 0.861764713 |
| fukami579 | 31249942 | 30673342 | 15336671 | 15095938 | 13669761 | 0.891312137 |
| fukami580 | 38602218 | 38100532 | 19050266 | 18659158 | 16669391 | 0.87502143 |
| fukami581 | 30560426 | 30008646 | 15004323 | 14738717 | 13103104 | 0.873288585 |
| fukami582 | 44642036 | 44176324 | 22088162 | 20580542 | 19024093 | 0.861280038 |
| fukami583 | 31568086 | 31007708 | 15503854 | 15065531 | 13690761 | 0.883055336 |
| fukami584 | 48685560 | 48399818 | 24199909 | 22590205 | 20855508 | 0.861801092 |
| fukami585 | 41781996 | 41613802 | 20806901 | 19359789 | 17781687 | 0.854605258 |
| fukami586 | 45288450 | 45015878 | 22507939 | 20948870 | 19219362 | 0.853892575 |
| fukami587 | 39453864 | 39017264 | 19508632 | 18152972 | 16655156 | 0.853732645 |
| fukami588 | 38634340 | 38316570 | 19158285 | 17787209 | 16346470 | 0.853232427 |
| fukami589 | 58464228 | 58036704 | 29018352 | 26999656 | 24661896 | 0.849872384 |
| fukami590 | 30504334 | 30021558 | 15010779 | 13824290 | 12624538 | 0.841031501 |
| fukami591 | 33405734 | 33062556 | 16531278 | 15333198 | 13960716 | 0.844503129 |
| fukami592 | 33121926 | 32960454 | 16480227 | 15344133 | 14098060 | 0.855453023 |
| fukami593 | 33365282 | 32853804 | 16426902 | 16112828 | 14542292 | 0.88527295 |
| fukami594 | 45094298 | 44683830 | 22341915 | 21833701 | 19740611 | 0.883568441 |
| fukami595 | 31643744 | 31255666 | 15627833 | 14488465 | 13382670 | 0.856335616 |
| fukami596 | 32638632 | 32077640 | 16038820 | 15768769 | 14349604 | 0.894679534 |
| fukami597 | 36991658 | 36295712 | 18147856 | 17777682 | 16190147 | 0.892124502 |
| fukami600 | 29048188 | 28802768 | 14401384 | 14063062 | 12749218 | 0.885277276 |
| fukami601 | 34483600 | 34000376 | 17000188 | 16741877 | 14964084 | 0.880230501 |
| fukami602 | 31741528 | 30985320 | 15492660 | 13378145 | 12396328 | 0.800142003 |
| fukami603 | 43932610 | 43507118 | 21753559 | 21320810 | 19581257 | 0.900140386 |
| fukami604 | 30292266 | 29727234 | 14863617 | 12882029 | 11931174 | 0.80271 |
| fukami605 | 32847814 | 32612298 | 16306149 | 15970865 | 14668313 | 0.899557155 |
| fukami606 | 32426294 | 32116592 | 16058296 | 15742121 | 14413128 | 0.897550276 |
| fukami607 | 23583516 | 23472978 | 11736489 | 11509356 | 10597225 | 0.902929743 |
| fukami608 | 39442566 | 39033188 | 19516594 | 19102717 | 17557810 | 0.899634947 |
| fukami609 | 58520278 | 57400426 | 28700213 | 24793446 | 23001214 | 0.80143008 |
| fukami610 | 24543274 | 19317256 | 9658628 | 9467202 | 8646302 | 0.895189462 |
| fukami611 | 34322142 | 33506384 | 16753192 | 16511884 | 14830412 | 0.885229036 |
| fukami612 | 29375032 | 29077118 | 14538559 | 14311636 | 12567831 | 0.864448189 |
| fukami613 | 40435872 | 40225742 | 20112871 | 19801100 | 17597308 | 0.874927702 |
| fukami614 | 38811054 | 38579656 | 19289828 | 18966171 | 16981806 | 0.880350307 |
| fukami615 | 41143316 | 40432664 | 20216332 | 19880879 | 17783476 | 0.879658882 |
| fukami616 | 44775492 | 44438628 | 22219314 | 21850977 | 19584866 | 0.881434323 |
| fukami617 | 38338306 | 38146676 | 19073338 | 17840951 | 16506962 | 0.86544694 |
| fukami618 | 40737880 | 40387716 | 20193858 | 18889808 | 17476803 | 0.865451416 |
| fukami619 | 46039728 | 45638322 | 22819161 | 21318660 | 19732072 | 0.864715052 |
| fukami620 | 33995758 | 33695814 | 16847907 | 15668899 | 14511927 | 0.86134895 |
| fukami621 | 33870982 | 33537378 | 16768689 | 15651934 | 14442613 | 0.861284564 |
| fukami622 | 33178548 | 31955992 | 15977996 | 15615557 | 14089117 | 0.881782484 |
| fukami623 | 42976510 | 42428292 | 21214146 | 19793825 | 18272665 | 0.861343417 |
| fukami624 | 29874174 | 28926054 | 14463027 | 12488771 | 11578141 | 0.800533733 |
| fukami625 | 36640766 | 36327180 | 18163590 | 17900470 | 16023740 | 0.88219014 |
| fukami626 | 34439332 | 33763460 | 16881730 | 16614887 | 14705631 | 0.871097393 |
| fukami627 | 41068546 | 40530698 | 20265349 | 19940129 | 17930501 | 0.884786193 |
| fukami628 | 29733862 | 29262980 | 14631490 | 14386254 | 12953399 | 0.88530963 |
| fukami629 | 36964058 | 36776132 | 18388066 | 18109695 | 16270695 | 0.884850805 |
| fukami630 | 30739320 | 29541456 | 14770728 | 14255362 | 12917895 | 0.874560482 |
| fukami631 | 39177710 | 38768506 | 19384253 | 19085617 | 17075367 | 0.880888575 |
| fukami632 | 48967564 | 48682074 | 24341037 | 23946459 | 21559407 | 0.885722617 |
| fukami633 | 39560240 | 39283162 | 19641581 | 19360241 | 17731379 | 0.902747034 |
| fukami634 | 51039712 | 50438584 | 25219292 | 24850741 | 22753286 | 0.902217477 |
| fukami635 | 30516198 | 29542590 | 14771295 | 13635635 | 12465339 | 0.843889381 |
| fukami637 | 38957466 | 37915806 | 18957903 | 17518102 | 15787063 | 0.832743105 |
| fukami638 | 35459940 | 33932722 | 16966361 | 16663813 | 14941335 | 0.880644647 |
| fukami639 | 42948172 | 42748100 | 21374050 | 21055686 | 19374664 | 0.906457316 |
| fukami640 | 36379372 | 35956958 | 17978479 | 17715783 | 16219809 | 0.902179155 |
| fukami641 | 26390594 | 26102660 | 13051330 | 12826659 | 11738217 | 0.899388568 |
| fukami642 | 35409958 | 34972708 | 17486354 | 17187638 | 15595303 | 0.891855615 |
| fukami643 | 57210412 | 56353746 | 28176873 | 27669415 | 25047901 | 0.888952475 |
| fukami644 | 33874146 | 32831724 | 16415862 | 16118570 | 14653322 | 0.892631895 |
| fukami645 | 43394718 | 43066330 | 21533165 | 21163147 | 19223799 | 0.892753063 |
| fukami646 | 41151892 | 40841246 | 20420623 | 20015446 | 18190979 | 0.890814105 |
| fukami647 | 34049394 | 33815614 | 16907807 | 16573305 | 15088403 | 0.892392668 |
| fukami648 | 34752876 | 34494284 | 17247142 | 16937156 | 15333205 | 0.889028745 |
| fukami649 | 51596786 | 50034338 | 25017169 | 22959211 | 21001728 | 0.83949259 |
| fukami650 | 39989122 | 39432148 | 19716074 | 16689418 | 15470032 | 0.784640593 |
| fukami651 | 31299514 | 30988326 | 15494163 | 13298883 | 12356830 | 0.797515167 |
| fukami652 | 25319068 | 25045098 | 12522549 | 12356287 | 11048987 | 0.882327312 |
| fukami653 | 21106790 | 16622366 | 8311183 | 7646497 | 7070471 | 0.850717762 |
| fukami654 | 31638996 | 31136632 | 15568316 | 15360422 | 13968619 | 0.897246626 |
| fukami655 | 48474524 | 46324418 | 23162209 | 22658650 | 20204864 | 0.872320252 |
| fukami656 | 26168612 | 25143404 | 12571702 | 10773005 | 9993756 | 0.794940574 |
| fukami657 | 30894610 | 30575180 | 15287590 | 15109097 | 13731572 | 0.89821692 |
| fukami658 | 33015726 | 32529132 | 16264566 | 15755748 | 14251810 | 0.876249019 |
| fukami659 | 59003224 | 58686728 | 29343364 | 28995665 | 26355903 | 0.898189553 |
| fukami660 | 39569988 | 39280610 | 19640305 | 19409068 | 17587412 | 0.895475503 |
| fukami661 | 37753304 | 37421316 | 18710658 | 18481132 | 16864271 | 0.90131897 |
| fukami662 | 33685912 | 33340642 | 16670321 | 16467103 | 15064282 | 0.903658784 |
| fukami663 | 42774382 | 42404216 | 21202108 | 20942318 | 19052789 | 0.898627108 |
| fukami664 | 26270042 | 25842538 | 12921269 | 11934208 | 10929348 | 0.845841689 |
| fukami665 | 38720290 | 38456856 | 19228428 | 18938802 | 17171220 | 0.893012159 |
| fukami667 | 46765660 | 46245462 | 23122731 | 22754278 | 20603954 | 0.891069225 |
| fukami668 | 38363166 | 38119812 | 19059906 | 18763159 | 16999684 | 0.891908071 |
| fukami669 | 38408712 | 37910346 | 18955173 | 18673524 | 16956258 | 0.894545146 |
| fukami670 | 36775110 | 36001322 | 18000661 | 17758575 | 16151755 | 0.897286772 |
| fukami671 | 32230770 | 31868360 | 15934180 | 15670237 | 14228962 | 0.892983636 |
| fukami672 | 40534486 | 40342826 | 20171413 | 19878610 | 18202741 | 0.902402871 |
| fukami681 | 22486986 | 22154828 | 11077414 | 10910949 | 9839874 | 0.88828259 |
| fukami682 | 37363160 | 36820558 | 18410279 | 17895156 | 16192336 | 0.879526921 |
| fukami683 | 70741788 | 69750038 | 34875019 | 34400940 | 31211960 | 0.894966107 |
| fukami684 | 45137006 | 44748066 | 22374033 | 22057140 | 20030440 | 0.895253887 |
| fukami685 | 46824446 | 46184212 | 23092106 | 22761206 | 20657025 | 0.894549202 |
| fukami686 | 51404688 | 51056740 | 25528370 | 25157278 | 22897053 | 0.896925773 |
| fukami687 | 36042450 | 35609724 | 17804862 | 17568356 | 16065132 | 0.902289049 |
| fukami688 | 28412368 | 28052382 | 14026191 | 13821990 | 12612010 | 0.899175692 |
| fukami689 | 35327372 | 34631052 | 17315526 | 14688920 | 13611271 | 0.786073204 |
| fukami690 | 43167854 | 42617588 | 21308794 | 20977634 | 19159142 | 0.899119021 |
| fukami691 | 36754048 | 36351312 | 18175656 | 17891242 | 16319810 | 0.897893864 |
| fukami692 | 35778778 | 35212576 | 17606288 | 17352943 | 15695331 | 0.891461676 |
| fukami693 | 43234742 | 42991136 | 21495568 | 21178909 | 19292162 | 0.89749487 |
| fukami694 | 28474380 | 28201314 | 14100657 | 13899872 | 12554796 | 0.890369576 |
| fukami696 | 37420712 | 37171038 | 18585519 | 18308250 | 16678072 | 0.897369183 |
| fukami129 | 29218560 | 28858424 | 14429212 | 14222734 | 12804787 | 0.887421087 |
| fukami130 | 35537604 | 35218724 | 17609362 | 16298761 | 14847686 | 0.843170014 |
| fukami131 | 26689960 | 26051784 | 13025892 | 11430574 | 10261082 | 0.787745054 |
| fukami132 | 22533060 | 21887634 | 10943817 | 9626187 | 8614797 | 0.787183941 |
| fukami133 | 34750938 | 34352124 | 17176062 | 14672752 | 13472869 | 0.784398019 |
| fukami134 | 26307050 | 25788954 | 12894477 | 11476939 | 10623854 | 0.823907321 |
| fukami135 | 26946604 | 26700220 | 13350110 | 11131510 | 9867804 | 0.739155258 |
| fukami136 | 29499006 | 28805878 | 14402939 | 12696678 | 11548073 | 0.801785872 |
| fukami290 | 56640686 | 56042994 | 28021497 | 27494772 | 24935662 | 0.889876155 |
| fukami81 | 32137394 | 31894876 | 15947438 | 14692833 | 13366384 | 0.838152436 |
| fukami82 | 25555362 | 25201260 | 12600630 | 11709744 | 10639991 | 0.84440151 |
| fukami83 | 24780454 | 24154080 | 12077040 | 11284085 | 10220248 | 0.84625438 |
| fukami84 | 44128828 | 43582668 | 21791334 | 20388870 | 18487714 | 0.848397533 |
| fukami85 | 36287540 | 35984334 | 17992167 | 16801628 | 15451426 | 0.858786271 |
| fukami86 | 21422594 | 21170762 | 10585381 | 10060569 | 9123869 | 0.861931092 |
| fukami87 | 41547828 | 41206382 | 20603191 | 19129876 | 17468314 | 0.847845074 |
| fukami88 | 55385854 | 55017646 | 27508823 | 25526476 | 23236842 | 0.844705061 |
| fukami673 | 50038262 | 49344412 | 24672206 | 24265840 | 22023073 | 0.892626829 |
| fukami674 | 35818756 | 34422836 | 17211418 | 16906540 | 15318388 | 0.89001313 |
| fukami676 | 3795330 | 3607522 | 1803761 | 1772087 | 1576584 | 0.874053713 |
| fukami677 | 36486990 | 35919428 | 17959714 | 17670839 | 15998954 | 0.890824542 |
| fukami678 | 45198544 | 44937166 | 22468583 | 22128635 | 20147652 | 0.896703277 |
| fukami679 | 25758902 | 25269720 | 12634860 | 11669059 | 10698151 | 0.846717019 |
| fukami680 | 19079832 | 18618020 | 9309010 | 8616789 | 7834536 | 0.841607862 |
| **Sum** | 25534966184 | 25118244274 | 12559122137 | 11776721540 | 10705747326 | 631.2864614 |
| **Average** | 34413701.06 | 33852081.23 | 16926040.62 | 15871592.37 | 14428230.9 | 0.8507903793 |

#### **S3: Top significantly differentially expressed genes**

| **Mean Expression** | **log2FoldChange** | **lfcSE** | **stat** | **p-value** | **padj** | **transcriptId** | **Signature description** | **start** | **stop** | **score** |
| --- | --- | --- | --- | --- | --- | --- | --- | --- | --- | --- |
| 1915.85783 | 0.54234786 | 0.11064073 | 4.90188273 | 9.49E-07 | 3.45E-05 | 105485 | Methyltransferase domain | 97 | 236 | 1.90E-07 |
| 7.52225423 | 0.6208208 | 0.43633254 | 1.42281572 | 0.15478958 | 0.28499523 | 112786 | - | 44 | 210 | 8.50E-06 |
| 264.870832 | -0.6428308 | 0.09581748 | -6.7089088 | 1.96E-11 | 1.02E-08 | 120870 | Ferredoxin reductase-type FAD binding domain profile. | 426 | 532 | 10.736 |
| 15.6372856 | -0.5398478 | 0.20859953 | -2.5879626 | 0.00965455 | 0.03686979 | 150265 | NAD dependent epimerase/dehydratase family | 5 | 232 | 5.70E-10 |
| 28.7370393 | 0.99149817 | 1.00477812 | 0.98678321 | 0.32374893 | 0.48534747 | 151961 | NA | NA | NA | NA |
| 23.2598683 | -0.958332 | 0.26842703 | -3.5701768 | 0.00035674 | 0.00299454 | 159807 | NA | NA | NA | NA |

#### **S4: Significantly enriched GO terms**

| [**GO.ID**](http://go.id) | **Term** | **Annotated Genes** | **Significant Genes** | **Expected # Genes** | **p-value** | **ontology** |
| --- | --- | --- | --- | --- | --- | --- |
| GO:0006096 | glycolytic process | 12 | 11 | 3.66 | 1.70E-05 | Biological Process |
| GO:0006099 | tricarboxylic acid cycle | 13 | 11 | 3.96 | 8.20E-05 | Biological Process |
| GO:0006412 | translation | 199 | 76 | 60.65 | 1.00E-04 | Biological Process |
| GO:0042364 | water-soluble vitamin biosynthetic process | 27 | 13 | 8.23 | 0.0041 | Biological Process |
| GO:0006575 | cellular modified amino acid metabolic process | 19 | 9 | 5.79 | 0.0041 | Biological Process |
| GO:0009064 | glutamine family amino acid metabolic process | 16 | 10 | 4.88 | 0.0079 | Biological Process |
| GO:1901566 | organonitrogen compound biosynthetic process | 428 | 165 | 130.44 | 0.0079 | Biological Process |
| GO:0019752 | carboxylic acid metabolic process | 185 | 77 | 56.38 | 0.01 | Biological Process |
| GO:0006518 | peptide metabolic process | 211 | 83 | 64.3 | 0.0101 | Biological Process |
| GO:0006066 | alcohol metabolic process | 24 | 12 | 7.31 | 0.0116 | Biological Process |
| GO:0044271 | cellular nitrogen compound biosynthetic process | 597 | 193 | 181.94 | 0.0128 | Biological Process |
| GO:1901565 | organonitrogen compound catabolic process | 82 | 18 | 24.99 | 0.0133 | Biological Process |
| GO:0098662 | inorganic cation transmembrane transport | 50 | 20 | 15.24 | 0.019 | Biological Process |
| GO:1901135 | carbohydrate derivative metabolic process | 157 | 64 | 47.85 | 0.0205 | Biological Process |
| GO:0044283 | small molecule biosynthetic process | 113 | 53 | 34.44 | 0.0208 | Biological Process |
| GO:0006364 | rRNA processing | 28 | 14 | 8.53 | 0.0232 | Biological Process |
| GO:0009117 | nucleotide metabolic process | 78 | 36 | 23.77 | 0.0271 | Biological Process |
| GO:0000278 | mitotic cell cycle | 27 | 14 | 8.23 | 0.0278 | Biological Process |
| GO:0030001 | metal ion transport | 34 | 16 | 10.36 | 0.0288 | Biological Process |
| GO:0006457 | protein folding | 34 | 16 | 10.36 | 0.03 | Biological Process |
| GO:0018130 | heterocycle biosynthetic process | 390 | 114 | 118.86 | 0.0305 | Biological Process |
| GO:1901137 | carbohydrate derivative biosynthetic process | 106 | 36 | 32.3 | 0.0341 | Biological Process |
| GO:1901362 | organic cyclic compound biosynthetic process | 403 | 120 | 122.82 | 0.0382 | Biological Process |
| GO:0003735 | structural constituent of ribosome | 131 | 63 | 41.07 | 3.20E-05 | Molecular Function |
| GO:0016616 | Catalysis of an oxidation-reduction (redox) reaction | 32 | 20 | 10.03 | 0.00026 | Molecular Function |
| GO:0051287 | NAD binding | 25 | 15 | 7.84 | 0.00279 | Molecular Function |
| GO:0008289 | lipid binding | 29 | 10 | 9.09 | 0.0049 | Molecular Function |
| GO:0043168 | anion binding | 542 | 178 | 169.91 | 0.00533 | Molecular Function |
| GO:0000287 | magnesium ion binding | 32 | 17 | 10.03 | 0.00823 | Molecular Function |
| GO:0022890 | inorganic cation transmembrane transport | 74 | 28 | 23.2 | 0.01403 | Molecular Function |
| GO:0019842 | vitamin binding | 51 | 18 | 15.99 | 0.01426 | Molecular Function |
| GO:0004519 | endonuclease activity | 30 | 14 | 9.4 | 0.03102 | Molecular Function |
| GO:1901363 | heterocyclic compound binding | 1021 | 329 | 320.07 | 0.03466 | Molecular Function |
| GO:0097159 | organic cyclic compound binding | 1021 | 329 | 320.07 | 0.03466 | Molecular Function |
| GO:0016667 | oxidoreductase activity, acting on a sulfur group of donors | 20 | 8 | 6.27 | 0.03483 | Molecular Function |
| GO:0016301 | kinase activity | 165 | 63 | 51.73 | 0.04357 | Molecular Function |
| GO:0005840 | ribosome | 126 | 59 | 37.99 | 1.10E-05 | Cellular Compartment |
| GO:0098798 | mitochondrial protein-containing complex | 36 | 11 | 10.85 | 0.009 | Cellular Compartment |
| GO:0016020 | membrane | 521 | 169 | 157.08 | 0.013 | Cellular Compartment |
| GO:0140534 | A protein complex that is part of an endoplasmic reticulum | 20 | 10 | 6.03 | 0.048 | Cellular Compartment |

#### **S5: eQTL results**

| **Signature description** | **Amino acid transporter** | **ABC transporter** |
| --- | --- | --- |
| Transcript ID | 38411 | 35539 |
| beta_G_ | -0.0573249 | 0.467644 |
| SE_G_ | 0.0177568 | 0.0659253 |
| Beta_E_ | -0.0692279 | 0.0636854 |
| SE_E_ | 0.0236893 | 0.0323827 |
| beta_GxE_ | 0.099928 | -0.36965 |
| SE_GxE_ | 0.0221908 | 0.0753043 |
| p_G_ (bonferroni) | 0.001 (2.08) | 3.33E-12 (0.049) |
| p_E_ (bonferroni) | 0.004 (5.72) | 0.049 (94.4) |
| p_GxE_ (bonferroni) | 7.9E-6 (0.013) | 1.15E-6 (0.002) |
| # effective tests | 1,594 | 1,901 |
